# Magnetogenetic cell activation using endogenous ferritin

**DOI:** 10.1101/2023.06.20.545120

**Authors:** Lisa Pomeranz, Rosemary Li, Xiaofei Yu, Leah Kelly, Gholamreza Hassanzadeh, Henrik Molina, Daniel Gross, Matthew Brier, George Vaisey, Putianqi Wang, Maria Jimenez-Gonzalez, Adolfo Garcia-Ocana, Jonathan Dordick, Jeffrey Friedman, Sarah Stanley

## Abstract

The ability to precisely control the activity of defined cell populations enables studies of their physiological roles and may provide therapeutic applications. While prior studies have shown that magnetic activation of ferritin-tagged ion channels allows cell-specific modulation of cellular activity, the large size of the constructs made the use of adeno-associated virus, AAV, the vector of choice for gene therapy, impractical. In addition, simple means for generating magnetic fields of sufficient strength have been lacking. Toward these ends, we first generated a novel anti-ferritin nanobody that when fused to transient receptor potential cation channel subfamily V member 1, TRPV1, enables direct binding of the channel to endogenous ferritin in mouse and human cells. This smaller construct can be delivered in a single AAV and we validated that it robustly enables magnetically induced cell activation *in vitro*. In parallel, we developed a simple benchtop electromagnet capable of gating the nanobody-tagged channel *in vivo*. Finally, we showed that delivering these new constructs by AAV to pancreatic beta cells in combination with the benchtop magnetic field delivery stimulates glucose-stimulated insulin release to improve glucose tolerance in mice *in vivo*. Together, the novel anti-ferritin nanobody, nanobody-TRPV1 construct and new hardware advance the utility of magnetogenetics in animals and potentially humans.

## Introduction

Technologies that allow for the temporal and spatial control of specific neuronal populations or circuits have enabled analyses of the functional effects of these neurons and may also have therapeutic applications(*1–3*). A range of tools exists, from optogenetics to chemically-gated receptors, each with their own features(*4*). Optogenetics enables the rapid alteration of neural activity but requires the implantation of an optical fiber(*5*). Chemogenetics does not require an implant but has slower onset kinetics(*6, 7*) and relies on the pharmacokinetics and pharmacodynamics of drugs and other chemical agents. Magnetic activation of ion channels potentially enables rapid, cell-specific neuromodulation without the need for implanted fibers(*8, 9*). Several independent groups have developed tools for magnetic modulation of neural activity and confirmed their efficacy in regulating neural activity(*9, 10*), calcium entry(*11–13*), cell migration(*14*), reward(*15*), metabolic control(*9*) and neural crest cell function(*8*).

Two technical requirements are necessary for deploying magnetogenetics. First, a genetically encoded version of an ion channel, such as a transient receptor potential (TRP) channel coupled to a ferritin nanoparticle must be delivered to specific cells. Second, a magnetic field of sufficient strength to gate the channel needs to be applied. In our initial studies, delivery of a magnetically susceptible channel was achieved by co-expressing green fluorescent protein (GFP)-labeled ferritin along with a TRPV1 fused to a commercially available anti-GFP nanobody(*9, 16*). The channel was then activated by placing freely moving animals in an MRI machine. Both of these steps had limitations. The delivery of both the channel and ferritin in the same vector required the use of a recombinant adenovirus which can carry a larger payload than adeno-associated virus (AAV), the vector of choice for animal and especially potential clinical studies. In addition, magnetic resonance imaging (MRI) machines are costly and access can be limited. In this report, we addressed the first limitation by generating an anti-ferritin nanobody that when fused to TRPV1 leads to direct binding of the channel to endogenous ferritin. The entire construct is only 2.9 kb and can be expressed in a single recombinant AAV. We then fabricated a simple electromagnet comprised of two antiparallel Helmholtz coils capable of delivering a sufficiently strong magnetic field to gate the channel. Finally, we validate the efficacy of the new construct *in vitro* and further demonstrated that the construct and this device can stimulate endogenous insulin release after AAV delivery *in vivo*. These new tools facilitate non-invasive magnetogenetic modulation of cell populations and in a companion publication, show that magnetogenetics can modulate neural activity *in vivo* and ameliorate the signs of Parkinson’s Disease in mice.

## Results

### Generation and validation of ferritin binding nanobodies

Magnetogenetic constructs rely on binding of multimodal TRP channels to ferritin. In previous work, we used a bicistronic construct to deliver a TRPV1-anti-GFP nanobody fusion protein together with GFP-tagged ferritin(*16*). However, these two constructs are 4.8 kb and increase to 5.5 kb when a cytomegalovirus (CMV) promoter is used to drive their expression. The maximal construct payload of an adeno-associated virus (AAV) is approximately 4.2 kb(*17*) and our previous construct is thus too large to deliver in a single AAV, the most commonly used viral vector in mice and humans. This approach also relied on the over-expression of an exogenously introduced GFP-tagged mouse ferritin, with unknown effects on cellular iron metabolism. To address these limitations, we set out to generate ferritin-binding nanobodies which offer high thermal stability, small size(*18*) and would potentially obviate the need for expressing GFP. The use of an anti-ferritin nanobody would also allow us to test whether binding to endogenous ferritin obviated the need for introducing the exogenous ferritin construct.

Llamas were immunized with human spleen ferritin every seven days for six weeks after which blood was collected for lymphocyte isolation (Fig 1A). In order to screen camelid nanobodies for ferritin binding using phage display technology, lymphocyte total RNA was extracted and used to generate cDNA followed by PCR amplification using variable heavy-chain antibody (VHH) encoding sequences and cloning into the phagemid vector, pMECS. A VHH library of ∼ 8x 10^7^ independent transformants was obtained. The library underwent three rounds of screening for binding to solid-phase coated human spleen ferritin (200 µg/ml, 20 μg/well) resulting in 8 x10^2^-fold enrichment in human ferritin-binding populations after the third round. In separate studies, the same library was also screened three times for binding to mouse liver ferritin (Gentaur, 200 µg/ml, 20 μg/well) leading to 5 x 10^2^-fold enrichment in mouse ferritin-binding populations after the third round of phage display. Periplasmic extracts from the 190 ferritin-binding phages that were isolated from the two screens were next analyzed for binding to human spleen ferritin and mouse liver ferritin by enzyme-linked immunosorbent assay (ELISA). In the first round ELISA assays, 133 of these 190 clones bound to human spleen ferritin (Fig S1A & B) and 54 of 190 clones showed significant binding to mouse ferritin (Fig S1C). Sequencing of the clones was performed and after removing duplicate clones, there were 59 unique clones from the screen for nanobodies to human ferritin and 19 unique clones from the screen for nanobodies to mouse ferritin.

**Figure 1:**
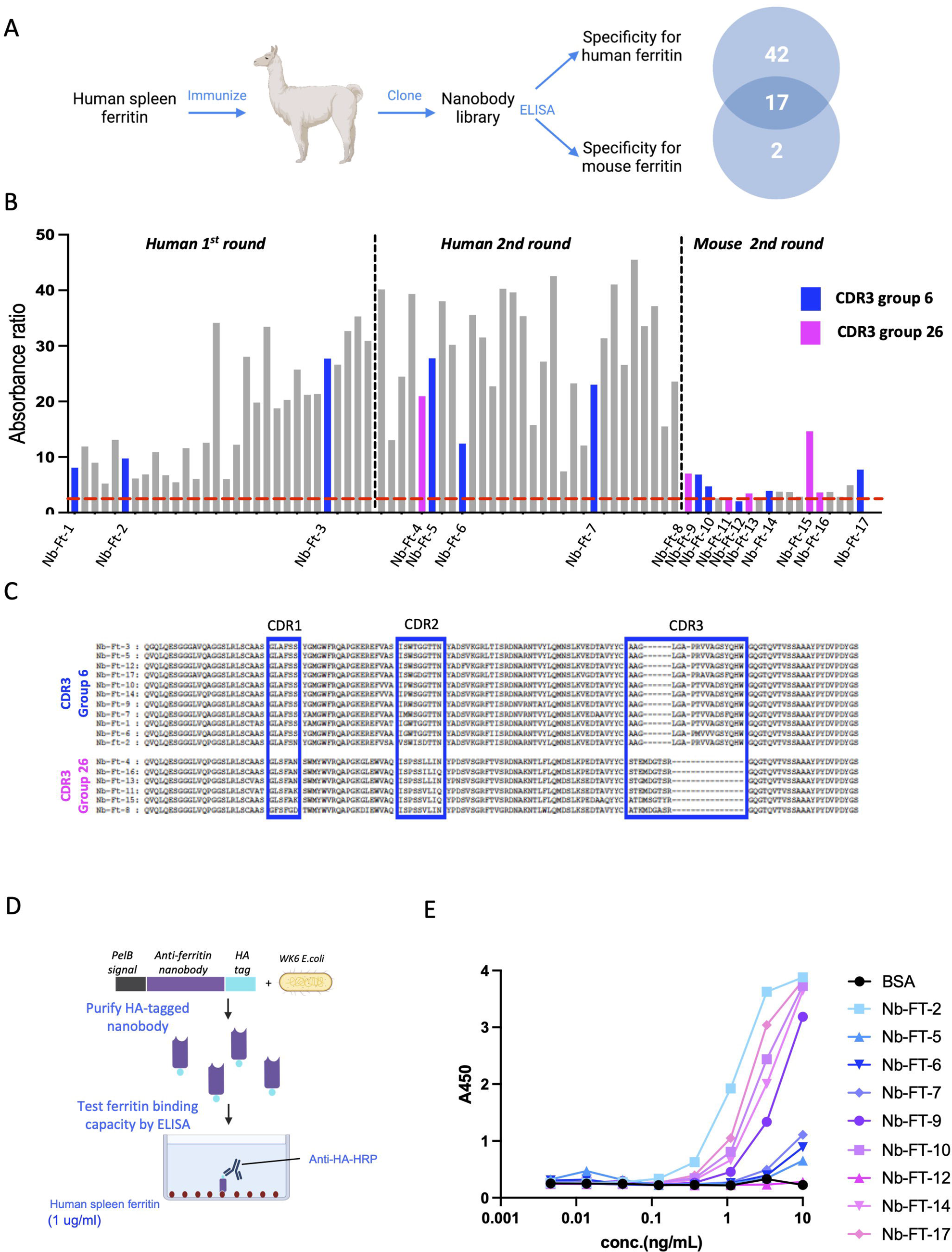
Generation of Nanobodies (Nb) to human and mouse ferritin and binding to human spleen ferritin. A. Schema of overall method for llama immunization, generation of phage display library, panning for selection of highly reactive nanobody clones to human spleen ferritin and mouse liver ferritin, ELISA and numbers of unique nanobodies reactive to human, mouse or both human and mouse ferritin. B. Example of ELISA selection for nanobodies from distinct groups for highly reactive clones. Clones were screened after the 1^st^ and 2^nd^ rounds of panning against human ferritin and after the 2^nd^ round of panning against mouse ferritin. Clones exhibiting absorbance ratio greater than 2.5 (dashed red line) were selected for further characterization. C. Alignment of amino acid sequence of nanobodies belonging to CDR3 groups 6 and 26 with the highest reactivity to human and mouse ferritin. CDR1, CDR2, and CDR3 are colored in blue. The nanobodies’ CDR3 group and names are on the right. D. Schema of method for quantification of nanobody binding to human spleen ferritin. HA-tagged nanobodies from CDR3 groups 6 and 26 were purified from transformed WK6 E. coli by metal-ion affinity chromatography. Serial dilutions of nanobodies or BSA (100ul) were incubated on plates coated with human spleen ferritin (1ug/ml) and binding quantified by ELISA after incubation with anti-HA-HRP antibody. E. Quantification of anti-ferritin nanobodies from CDR3 groups 6 and 26 binding to human spleen ferritin.

The nanobody clones were further classified based on the similarity of their complementarity determining region 3 (CDR3)(*19*) (Fig S1D). Sequence data demonstrated that the 59 human ferritin nanobodies belonged to 22 different CDR3 groups and that the 19 nanobodies binding mouse ferritin belonged to six different CDR3 groups. The 17 nanobodies from CDR3 groups 6 and 26 included nanobody clones that bound to both human ferritin (nanobody-Ferritin (Nb-Ft) clones 1-7) and mouse ferritin (Nb-Ft clones 8 - 17) (Figure 1B and 1C). Because we had initially set out to identify nanobodies capable of binding to both human and mouse ferritin these 17 nanobodies were characterized further.

We measured the affinity of the 17 nanobody clones belonging to CDR3 groups 6 and 26 to identify those that bound human ferritin most tightly. The nanobodies were fused to hemagglutinin (HA) and soluble protein was expressed in E. coli WK6 cells and purified by immobilized metal-ion affinity chromatography (Fig 1D). Binding of the partially purified nanobodies (0.0045 - 10ng/ml) to human spleen ferritin was analyzed by ELISA (Fig 1E). Five nanobody clones bound human ferritin with higher affinity (Half maximal binding: 1.2 ng/ml for clone 2, 1.9 ng/ml for clone 17, 2.8 ng/ml for clone 10, 3.9 ng/ml for clone 14 and 5.6 ng/ml for clone 9) (Fig 1E). We selected the five nanobodies with the highest affinity for human ferritin and quantitated binding to mouse ferritin. Binding to GFP-tagged mouse ferritin (GFP-mFerritin) was evaluated by tagging nanobodies with HA-mCherry and co-transfecting with a GFP-mFerritin construct (Fig 2A). As a control, we also tested the binding of GFP-mFerritin to the previously reported anti-GFP nanobody(*9*) that was also fused to HA-mCherry. The tagged nanobodies and their bound proteins were immunoprecipitated using anti-HA agarose beads and then analyzed using SDS-PAGE and mass spectrometry. SDS-PAGE analysis of the immunoprecipitated proteins from cells transfected with the anti-ferritin nanobodies showed a band of approximately 71 kDa corresponding to GFP-mFerritin and approximately 42 kDa corresponding to the HA-mCherry tagged-nanobodies (Fig 2B). Bands of the same 71 kDa size migrating with the same apparent molecular weight as ferritin were also seen after immunoprecipitations of lysates from cells that were transfected with the anti-GFP nanobody (lane 7). Binding of anti-ferritin nanobodies to ferritin was confirmed by mass spectrometry of immunoprecipitates. Endogenous human ferritin was highly enriched in the immunoprecipitates from HEK-293T cells expressing the anti-ferritin nanobody clones together with the transfected GFP-tagged mouse ferritin compared to HEK-293T cells expressing GFP-tagged mouse ferritin alone (Enrichment in human ferritin: Nb-Ft clone 2: 58.1-fold enrichment, clone 9: 4.6, clone 10: 7.0, clone 14: 5.1, clone 17: 20.9) (Fig 2C). In addition, in HEK-293T cells expressing the GFP-tagged mouse ferritin constructs, immunoprecipitates from cells expressing anti-ferritin nanobody clones 2 and 17 or an anti-GFP nanobody showed highly significant enrichment for mouse ferritin compared to HEK-293T cells expressing GFP-tagged mouse ferritin alone (Enrichment in mouse ferritin: Nb-Ft clone 2: 14.5-fold enrichment, clone 9: 1.0 clone 10: 1.1, clone 14: 0.9, clone 17: 6.0 and anti-GFP nanobody: 44.3) (Fig 2C). Interestingly, human ferritin was enriched in immunoprecipitates from HEK-293T cells transfected with anti-GFP nanobody and GFP-tagged mouse ferritin suggesting that endogenous human ferritin subunits may assemble with GFP-tagged mouse ferritin subunits which can be immunoprecipitated with the anti-GFP nanobody. These findings show affinity of the anti-ferritin nanobodies for human and mouse ferritin. Having confirmed binding of the nanobodies to ferritin, we next evaluated their ability to activate a Ca^2+^ dependent reporter in a magnetic field.

**Figure 2:**
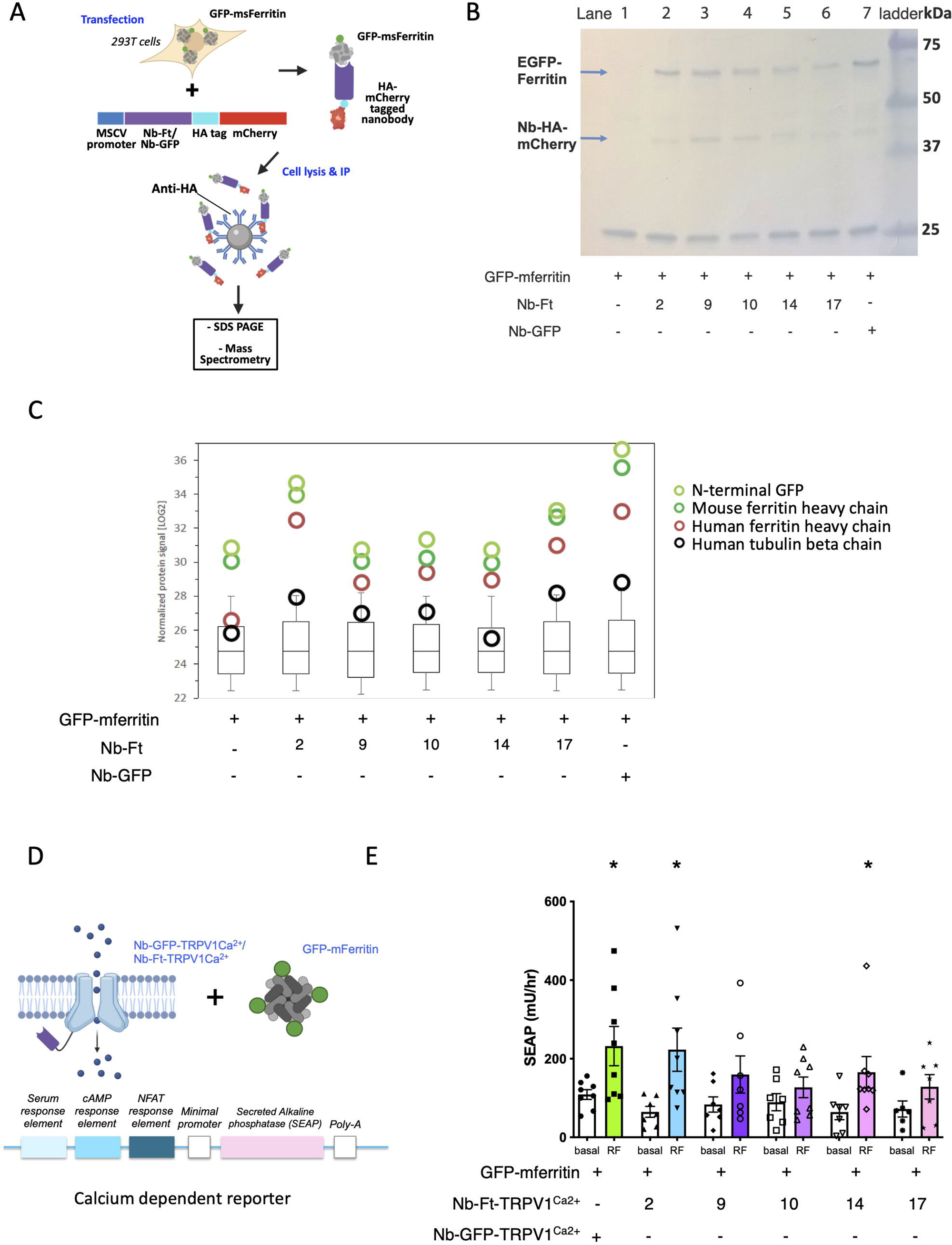
Screen of Nb clones to mouse ferritin. A. Schema of method to characterize Nb-Ft binding to mouse ferritin. Mammalian expression plasmids with Nb-Ft sequences (Nb-Ft-2, -9, -10, -14, -17) or Nb-GFP fused to HA-tagged mCherry were co-transfected into HEK-293T cells along with a plasmid expressing GFP-tagged mouse ferritin (GFP-mFerritin). After 48 hours, cells were lysed and HA-tagged nanobodies along with their bound proteins were immunoprecipitated using anti-HA conjugated agarose beads. Eluted proteins were examined by SDS-PAGE and mass spectrometry. B. SDS–PAGE analysis of immunoprecipitated Nb-Ft-HA-mCherry and bound GFP-tagged ferritin demonstrating bands of the expected sizes (42 kDa and 71 kDa respectively). Molecular weight markers, size indicated in kDa. C. Quantification of nanobody-bound proteins by mass spectrometry normalized for protein signal (log2). The protein signals for mouse ferritin heavy chain (transfected as GFP-mFerritin), human ferritin heavy chain (FTH1_human, endogenous to HEK-293T cells), and N-terminal GFP moiety of GFP-mFerritin are shown along with a control peptide, human tubulin (TBB5_human). The overall protein signal shown in the box plot does not differ between test conditions. D. Schema of method to validate ability of Nb-ferritin clones to transduce magnetic field into cell activation. Constructs with Nb-Ft sequences (Nb-Ft-2, -9, -10, -14, -17) fused to the N-terminal of TRPV1^Ca2+^ were generated and transfected into HEK-293T cells along with a plasmid expressing GFP-tagged mouse ferritin (GFP-mFerritin) and a calcium-dependent secreted alkaline phosphatase (SEAP). Cells transfected Nb-GFP-TRPV1^Ca2+^/GFP-mFerritin and the reporter were used as a positive control. E. Oscillating magnetic field treatment (465kHz, 30mT) significantly increases calcium-dependent SEAP release from 293T cells transfected with Nb-GFP-TRPV1^Ca2+^, Nb-Ft-2-TRPV1^Ca2+^ and Nb-Ft-14-TRPV1^Ca2+^ and GFP-mFerritin. Data analyzed by two-tailed, unpaired t-test with Welch’s correction. Nb-GFP-TRPV1^Ca2+^, basal vs. RF * p = 0.04, n = 8 & 8; Nb-Ft-2-TRPV1^Ca2+^, basal vs. RF * p = 0.02, n = 7 & 8; Nb-Ft-14-TRPV1^Ca2+^, basal vs. RF * p = 0.04, n = 7 & 8.

We generated five separate constructs in which the anti-ferritin nanobody clones 2, 9, 10, 14 and 17 were fused to the N-terminus of TRPV1. These were then cloned into bicistronic constructs in which a 2A peptide cleavage site was followed by GFP-tagged mouse ferritin in the pMSCV mammalian expression vector (pMSCV-Nb-Ft – TRPV1^Ca2+^-2A-GFP-mFerritin). Plasmids for each of the five nanobody-TRPV1 fusions were co-transfected into HEK-293T cells together with a calcium-dependent secreted alkaline phosphatase (SEAP) reporter construct (Fig 2D). The promoter in this reporter, which has been previously published(*20, 21*), includes three Ca2+ response elements in cis (serum response element, cyclic adenosine monophosphate response element, and nuclear factor of activated T cell response element) upstream of a minimal promoter. This promoter is activated after gating of TRPV1 and controls the synthesis of SEAP(*20*) providing a cumulative indicator of Ca^2+^ entry into cells after exposure to a magnetic field. Exposure of cells expressing the TRPV1 nanobody fusions and GFP-ferritin to an oscillating magnetic field (465kHz, 30mT) resulted in a significant increase of SEAP production from cells transfected with anti-ferritin nanobodies 2 and 14 fused to TRPV1 compared to transfected cells without magnet treatment (basal)(Fig 2E, Nb-Ft-TRPV1^Ca2+^ clone 2 -Basal 64.7 ± 13.9 mU/hr, Magnet: 223.0 ± 55.0 mU/hr, p < 0.05, Nb-Ft-TRPV1^Ca2+^ clone 14 -Basal 64.3 ± 19.4 mU/hr, Magnet: 165.5 ± 19.4 mU/hr, p < 0.05). Ferritin nanobody stimulation of SEAP production was compared with a previously characterized construct (pMSCV-Nb-GFP – TRPV1^Ca2+^-2A-GFP-mFerritin)(*16*) in which TRPV1 is fused to an anti-GFP nanobody that tethers the transfected GFP-tagged ferritin to the channel (Nb-GFP-TRPV1^Ca2+^ -basal 108.9 ± 12.4 mU/hr, Magnet: 232.2 ± 49.8 mU/hr, p < 0.05). The magnitude of the response was similar in cells expressing the TRPV1-anti-ferritin nanobody fusions for clones 2, 9 and 14 and those expressing the TRPV1 anti-GFP nanobody fusion.

Together, these findings confirm that the newly generated anti-ferritin nanobodies bind both human and mouse ferritin and can tether TRPV1 to ferritin to enable magnetogenetic activation. The aggregate data further revealed that anti-ferritin nanobody clone 2 had the highest affinity for human ferritin, the greatest enrichment of mouse ferritin after immunoprecipitation and showed the most significant increase of calcium-dependent SEAP in a magnetic field. Thus, anti-ferritin nanobody clone 2 was selected for further studies.

### Validation of anti-ferritin clone 2 for magnet stimulation of cells in vitro

The binding affinity of anti-ferritin nanobody clone 2 to human ferritin in solution was calculated using isothermal titration calorimetry and showed an equilibrium dissociation of 0.54 ± 0.2 μM (Fig 3A and B). We next confirmed that TRPV1 fused to anti-ferritin nanobody clone 2 (referred to hereafter as Nb-Ft-2-TRPV1^Ca2+^) could activate cells using a second calcium dependent reporter assay and also tested its ability to directly gate calcium in a magnetic field. HEK-293T cells expressing Nb-Ft-2-TRPV1^Ca2+^, and GFP-mFerritin along with the same calcium-responsive promoter described above (Fig 2D) cloned upstream of furin-modified insulin(*21*) were placed in magnetic field. Exposure of HEK-293T cells transfected with pMSCV-Nb-Ft-2-TRPV1^Ca2+^-T2A-GFP-mFerritin and pSRE-CRE-NFAT-insulin to a magnetic field (465kHz, 30mT) increased insulin secretion more than two-fold (Basal: 59.7 ± 12.6 µg/L, Magnet: 136.8 ± 13.9 µg/L, p < 0.001) and was in line with the effects of magnetic field treatment on insulin release in HEK-293T cells expressing the previously published Nb-GFP-TRPV1^Ca2+^ and GFP-mFerritin constructs (Basal: 21.3 ± 5.5 µg/L, Magnet: 67.6 ± 13.3 µg/L, p < 0.01) (Fig 3C).

**Figure 3:**
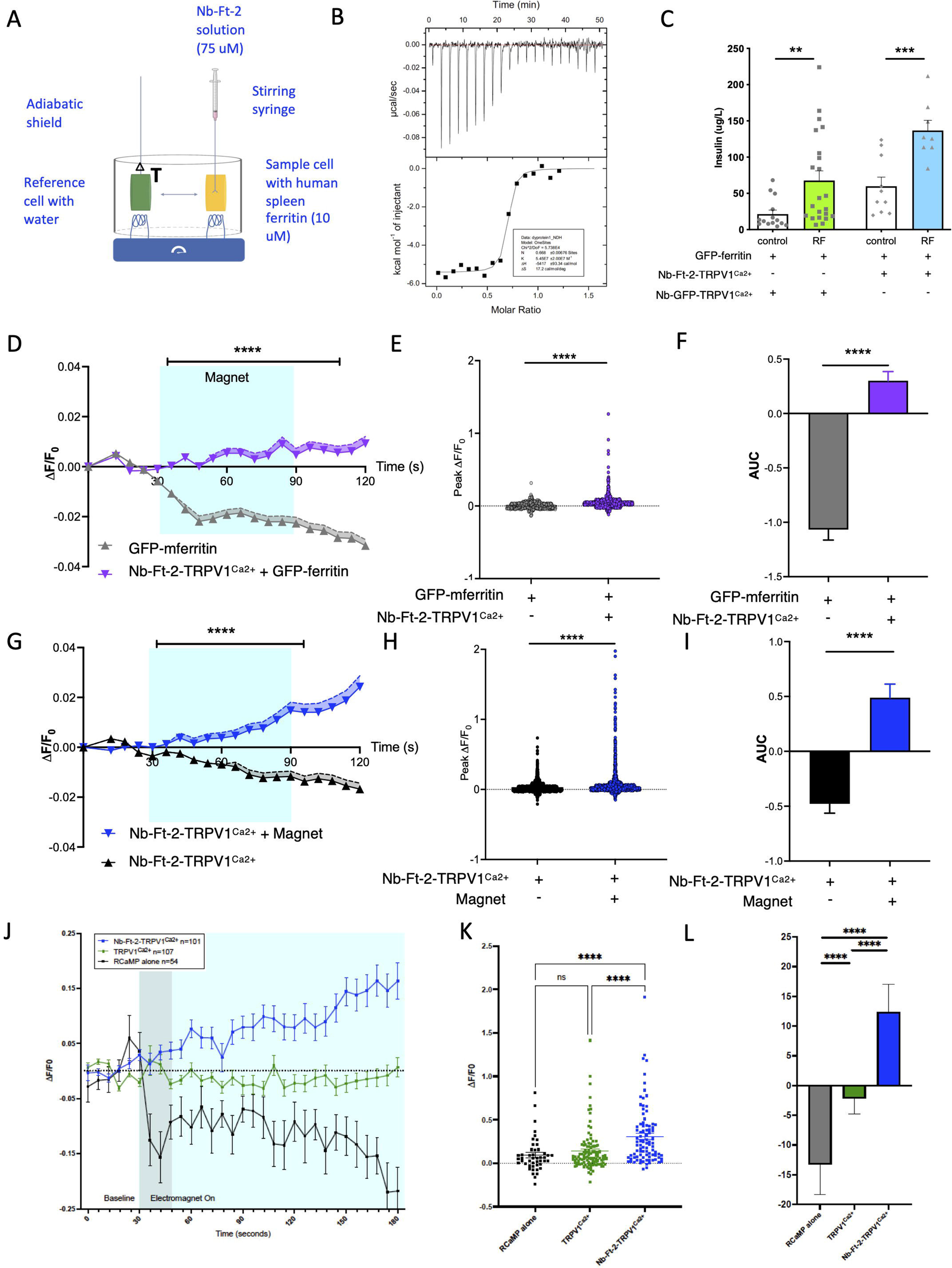
Functional validation of Nb-Ferritin clones *in vitro*. A. Schema of isothermal titration calorimetry assay for binding of anti-ferritin nanobody-2 (Nb-Ft2) to human spleen ferritin. Human spleen ferritin (10 μM) was titrated against 75 μM Nb-Ft2 (75 μM, single 0.4 μL injection, then 19 injections of 2 μL each, 150-sec intervals, 750 rpm stirring) with a reference power of 10 μcal/sec. B. Binding affinity and interaction between Nb-Ft2 and human spleen ferritin by isothermal titration calorimetry at 25°C (K_d_ = 0.54 ± 0.2 μM) .C. Oscillating magnetic field treatment (465kHz, 30mT) significantly increases calcium-dependent insulin release from 293T cells transfected with Nb-GFP-TRPV1^Ca2+^/GFP-mFerritin or with Nb-Ft-2-TRPV1^Ca2+^/GFP-mFerritin. Nb-GFP-TRPV1^Ca2+^/GFP-mFerritin: Data analyzed by Mann-Whitney test, basal vs. RF ** p = 0.003, n = 14 & 22. Nb-Ft-2-TRPV1^Ca2+^: Data analyzed by unpaired two-tailed t-test with Welch’s correction, basal vs. RF *** p = 0.0009, n = 10 & 8. D. Magnet treatment significantly increases normalized Fluo-4 fluorescence (DF/F_0_) in Neuro2A cells expressing Nb-Ft-2-TRPV1^Ca2+^/GFP-mFerritin (1627 cells) compared to cells expressing GFP-mFerritin alone (504 cells). Data were analyzed by two-way ANOVA with Tukey’s multiple comparison test, **** p < 0.0001 E. Peak ΔF/F_0_ with magnet treatment of Neuro2A cells expressing Nb-Ft-2-TRPV1^Ca2+^/GFP-mFerritin (1627 cells) or GFP-mFerritin alone (504 cells). Data were analyzed by Mann Whitney U test **** p < 0.0001. F. Cumulative change in ΔF/F_0_ with magnet treatment of Neuro2A cells expressing Nb-Ft-2-TRPV1^Ca2+^/GFP-mFerritin (1627 cells) or GFP-mFerritin alone (504 cells). Data were analyzed by Mann Whitney U test **** p < 0.0001. G. Normalized Fluo-4 fluorescence (ΔF/F_0_) in Neuro2A cells expressing Nb-Ft-2-TRPV1^Ca2+^ with (786 cells) or without (1659 cells) magnet treatment. Data were analyzed by two-way ANOVA with Tukey’s multiple comparison test, **** p < 0.0001 H. Peak ΔF/F_0_ of Neuro2A cells expressing Nb-Ft-2-TRPV1^Ca2+^ with (786 cells) or without (1659 cells) magnet treatment. Data were analyzed by Mann Whitney U test **** p < 0.0001. I. Cumulative change in ΔF/F_0_ of Neuro2A cells expressing Nb-Ft-2-TRPV1^Ca2+^ with (786 cells) or without (1659 cells) magnet treatment. Data were analyzed by Mann Whitney U test **** p < 0.0001. J. Changes in RCaMP fluorescence normalized to baseline fluorescence (ΔF/F_0_) with magnet treatment of HEK-293T cells expressing RCaMP alone (54 cells), TRPV1^Ca2+^ (107 cells) or Nb-Ft-2-TRPV1^Ca2+^ (101 cells). Error bars represent mean +/-SEM. K. Peak ΔF/F_0_ with magnet treatment of HEK-293T cells expressing RCaMP alone (54 cells), TRPV1^Ca2+^ (107 cells) or Nb-Ft-2-TRPV1^Ca2+^ (101 cells). Data were analyzed by ordinary one-way ANOVA with Tukey’s multiple comparison test, **** p < 0.0001. Error bars represent mean ± SD. L. Cumulative change in ΔF/F_0_ with magnet treatment of HEK-293T cells expressing RCaMP alone (54 cells), TRPV1^Ca2+^ (107 cells) or Nb-Ft-2-TRPV1^Ca2+^ (101 cells). AUC was calculated for the period of magnet exposure between 48-180 seconds (after focus adjustment) and analyzed by ordinary one-way ANOVA with Tukey’s multiple comparison test, **** p < 0.0001. Data are shown as mean ± SD.

We next generated an expression vector with Nb-Ft2 fused to TRPV1^Ca2+^ under the control of a neuronal human synapsin promoter (hSyn-Nb-Ft-2-TRPV1^Ca2+^). We confirmed cell surface expression of Nb-Ft-2-TRPV1^Ca2+^ by immunolabeling using antibodies directed to the extracellular portion of TRPV1 in transfected human HEK-293T and murine Neuro2A cells without permeabilization (Fig S2A and B). We also evaluated channel function in the membrane of cells expressing Nb-Ft-2-TRPV1^Ca2+^ by showing the appearance of capsaicin-dependent currents in pulled patches from Neuro2A cells transfected with this construct (Fig S2C). Capsaicin-induced currents were not observed in untransfected Neuro2A cells. The capsaicin-induced currents in Nb-Ft-2-TRPV1^Ca2+^ expressing Neuro2A cells also revealed the expected IV relationship for TRPV1(*22*). These studies confirm that the Nb-Ft-2-TRPV1^Ca2+^ fusion protein is incorporated into the cell membrane and that the TRPV1 channel is functional.

To further confirm the ability of hSyn-Nb-Ft-2-TRPV1^Ca2+^ to modulate cell activity in response to a magnetic field, we transfected murine Neuro2A cells with GFP-mFerritin, with and without hSyn-Nb-Ft-2-TRPV1^Ca2+^ then exposed the cells to a static magnetic field (190 mT) in the presence Fluo-4. Ca^2+^ levels in cells were quantified as the change in cell fluorescence divided by baseline fluorescence (ΔF/F0) (Fig 3D), peak ΔF/F0 (Fig 3E) and area under the curve (AUC) (Fig 3F). Magnetic exposure of Neuro2A cells expressing pAAV-hSyn-Nb-Ft-2-TRPV1^Ca2+^/GFP-mFerritin significantly increased peak ΔF/F0 and AUC compared to magnet treatment of cells expressing of GFP-mFerritin alone where imaging bleached the fluorescent signal over time (Peak ΔF/F0: 0.05 ± 0.003 for Nb-Ft-2-TRPV1/GFP-mFerritin, 0.004 ± 0.001 for GFP-mFerritin, p < 0.001. AUC: 0.3 ± 0.08 for Nb-Ft-2-TRPV1/GFP-mFerritin, −1.07 ± 0.09 GFP-mFerritin, p < 0.001).

Nb-Ft-2 binds to both mouse and human ferritin and we next considered the possibility that it might also tether endogenous ferritin thus obviating the need for co-transfection of GFP-mFerritin. Neuro2A cells expressing Nb-Ft-2-TRPV1^Ca2+^ alone (i.e.; without GFP-mFerritin) were exposed to the same magnetic field (190 mT) and intracellular calcium dynamics were measured using Fluo-4. The Ca^2+^ signal as measured by ΔF/F0, peak ΔF/F0 and AUC (Fig 3G, H, I) were all significantly increased in cells expressing Nb-Ft-2-TRPV1^Ca2+^ exposed to the magnetic field compared to cells expressing Nb-Ft-2-TRPV1^Ca2+^ without exposure to a magnetic field (Peak ΔF/F0: 0.07 ± 0.005 for Nb-Ft-2-TRPV1 + magnet, 0.02 ± 0.001 Nb-Ft-2-TRPV1, no magnet, p < 0.0001. AUC: 0.49 ± 0.12 Nb-Ft-2-TRPV1 + magnet, −0.47 ± 0.08 Nb-Ft-2-TRPV1, no magnet, p < 0.0001). Similarly, in human HEK-293T cells expressing Nb-Ft-2-TRPV1^Ca2+^, an oscillating magnetic field (465kHz, 30mT) significantly increased ΔF/F0 and peak ΔF/F0 RCaMP fluorescence compared to cells expressing wild type TRPV1^Ca2+^ without a Nb-Ft-2 fusion or expressing the reporter alone without transfection of TRPV1 (Fig 3J, K, L) (Peak ΔF/F0: 0.10 ± 0.027 for Control, 0.11 ± 0.019 for WT-TRPV1, 0.28 ± 0.034 for Nb-Ft-2-TRPV1, p < 0.0001 Kruskal-Wallis ANOVA. AUC: −13.27 ± 5.05 for Control, −2.19 ± 2.61 for WT-TRPV1, 12.38 ± 4.61 for Nb-Ft-2-TRPV1, *p=0.0195, ****p < 0.0001, Kruskal-Wallis). In contrast to the results of magnetic stimulation, fluorescence intensity still increased after capsaicin treatment of HEK-293T cells expressing wild type TRPV1 (Fig S2E & F). These data indicate that a magnetic field can increase intracellular calcium in mouse and human cells expressing Nb-Ft-2-TRPV1^Ca2+^ even without co-expression of the exogenous GFP-mFerritin construct. The Nb-Ft-2-TRPV1^Ca2++^ construct is 2.9 kb and these data suggested that magnogenetic activation could be achieved by introducing this construct into cells using an AAV that has a limited payload. We next tested this *in vitro* and *in vivo*.

### Magnet fields can stimulate insulin release *ex vivo* and *in vivo*

Calcium currents regulate the activity of many cell types and because magnetogenetic actuation does not require insertion of an implant, it is particularly well suited to modulating cell populations that are anatomically dispersed such as in the pancreas. We thus examined whether Nb-Ft-2-TRPV1^Ca2+^ could regulate the activity of pancreatic beta cells in culture and *in vivo*. To assess this, we generated an AAV construct using the small ubiquitously expressed JeT promoter to drive the cre-dependent expression of Nb-Ft-2-TRPV1^Ca2+^ with a double-floxed inverse open reading frame (DIO) (Fig 4A). This AAV construct was injected into multiple locations in the pancreas of Ins1-cre mice(*23*). Immunohistochemistry revealed TRPV1 expression in 8% of beta cells per mouse after four weeks as demonstrated by co-staining for insulin and an HA tag cloned between the Nb-Ft-2 and TRPV1 (Fig 4B). There was no detectable expression in alpha or delta cells in islets, assessed using antibodies to glucagon and somatostatin, or pancreatic tissue outside the islet (Fig S3A).

**Figure 4:**
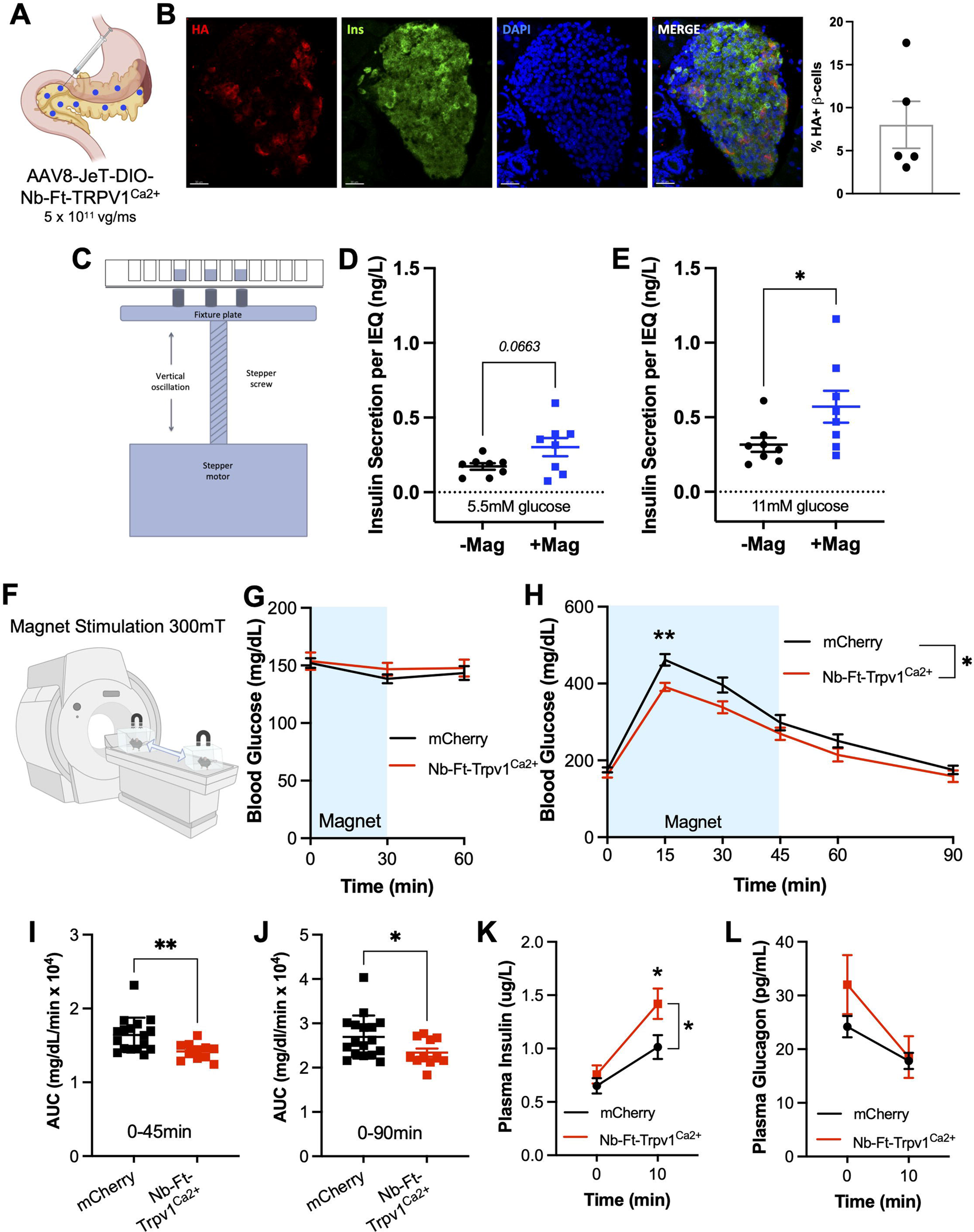
Efficacy of Nb-Ft-2-TRPV1^Ca2+^ *ex vivo* and *in vivo* in Ins1-cre mice. A. Schema of intrapancreatic injection of AAV8-JeT-DIO-Nb-Ft-2-TRPV1^Ca2+^ in Ins1-cre mice. B. Representative immunofluorescence images of islets from Ins1-cre mice after intrapancreatic injection of AAV8-JeT-DIO-Nb-Ft-2-TRPV1^Ca2+^ (HA-tagged Nb-Ft-2-TRPV1^Ca2+^(HA) in red, insulin (Ins) in green and DAPI in blue). Scale bars: 200µm. Right: Quantification of HA+ expression as a percentage insulin+ beta cells (n = 5 Ins1-cre mice). C. Schema of oscillating permanent magnet array for magnet treatment of islets *ex vivo*. D. Effects of oscillating magnet treatment on insulin secretion (normalized to islet equivalents, IEQ) at 5.5mM glucose from Ins1-cre mice injected with AAV8-JeT-DIO-Nb-Ft-2-TRPV1^Ca2+^. Data were analyzed by unpaired, two tail t-test, p = 0.066 E. Effects of oscillating magnet treatment on insulin secretion (normalized to islet equivalents, IEQ) at 11mM glucose from Ins1-cre mice injected with AAV8-JeT-DIO-Nb-Ft-2-TRPV1^Ca2+^. Data were analyzed by unpaired, two tail t-test, * p = 0.047 F. Schema of magnet stimulation of Ins1-cre mice with intrapancreatic injection of AAV8-JeT-DIO-Nb-Ft-2-TRPV1^Ca2+^ or AAV8-JeT-DIO-mCherry. G. Effects of magnet treatment on baseline blood glucose in Ins1-cre mice with intrapancreatic injection of AAV8-JeT-DIO-Nb-Ft-2-TRPV1^Ca2+^ or AAV8-JeT-DIO-mCherry. Data were analyzed by two-way ANOVA H. Effects of magnet treatment on blood glucose during glucose tolerance test (2g/kg) in Ins1-cre mice with intrapancreatic injection of AAV8-JeT-DIO-Nb-Ft-2-TRPV1^Ca2+^ or AAV8-JeT-DIO-mCherry. Data were analyzed by two-way repeated measures ANOVA with Sidak’s multiple comparison test. * p = 0.03 Nb-Ft-2-TRPV1^Ca2+^ vs. mCherry. ** p = 0.005 at 15 mins, Nb-Ft-2-TRPV1^Ca2+^ vs. mCherry. I. Effects of magnet treatment on cumulative blood glucose during 0-45 mins of glucose tolerance testing (2g/kg) in Ins1-cre mice with intrapancreatic injection of AAV8-JeT-DIO-Nb-Ft-2-TRPV1^Ca2+^ or AAV8-JeT-DIO-mCherry. Data were analyzed by unpaired two-tailed t-test. ** p = 0.004 Nb-Ft-2-TRPV1^Ca2+^ vs. mCherry. J. Effects of magnet treatment on cumulative blood glucose during 0-90 mins of glucose tolerance testing (2g/kg) in Ins1-cre mice with intrapancreatic injection of AAV8-JeT-DIO-Nb-Ft-2-TRPV1^Ca2+^ or AAV8-JeT-DIO-mCherry. Data were analyzed by unpaired two-tailed t-test. * p = 0.02 Nb-Ft-2-TRPV1^Ca2+^ vs. mCherry. K. Effects of magnet treatment on plasma insulin at 0 and 10 mins during a glucose tolerance test (2g/kg) in Ins1-cre mice with intrapancreatic injection of AAV8-JeT-DIO-Nb-Ft-2-TRPV1^Ca2+^ or AAV8-JeT-DIO-mCherry. Data were analyzed by two-way repeated measures ANOVA with Bonferroni’s multiple comparison test * p = 0.04 Nb-Ft-2-TRPV1^Ca2+^ vs. mCherry. * p = 0.018 Nb-Ft-2-TRPV1^Ca2+^ vs. mCherry at 10 mins. L. Effects of magnet treatment on plasma glucagon at 0 and 10 mins during a glucose tolerance test (2g/kg) in Ins1-cre mice with intrapancreatic injection of AAV8-JeT-DIO-Nb-Ft-2-TRPV1^Ca2+^ or AAV8-JeT-DIO-mCherry. Data were analyzed by two-way repeated measures ANOVA with Bonferroni’s multiple comparison test. p = 0.3 Nb-Ft-2-TRPV1^Ca2+^ vs. mCherry. Data are shown as mean ± SEM. AAV8-JeT-DIO-Nb-Ft-2-TRPV1^Ca2+^ n = 11, AAV8-JeT-DIO-mCherry n = 17.

We next isolated islets from the Ins1-cre mice, four weeks after injection of the AAV expressing the DIO-Nb-Ft-2-TRPV1^Ca2+^ construct or a control, DIO-mCherry construct and placed the cells in a cell culture incubator with a custom device for delivering an intermittent magnetic field. In this device, a permanent magnet array was placed immediately below the culture plate to expose the cells to a 120mT field. Movement of the plate 1 cm downward reduced the field strength to near 0 mT (Fig 4C). This cycle was repeated at 1 Hz with the displacement of the permanent magnet array controlled by a customized linear actuator and by IDEA® software (Haydon Kerk IDEA® Drive Interface Program) (Fig S3B). Exposure of the cultured islets isolated from Ins1-cre mice after AAV injections of DIO-Nb-Ft-2-TRPV1^Ca2+^ to an oscillating magnetic field (120mT, 1 Hz) significantly increased glucose-stimulated insulin secretion in the presence of 11mM glucose (Basal: 0.31 ± 0.05 ug/L per islet equivalent, Magnet: 0.57 ± 0.1 ug/L per islet equivalent, p < 0.05) with a trend to increased insulin secretion at 5.5mM glucose (Basal: 0.17 ± 0.02 ug/L per islet equivalent, Magnet: 0.30 ± 0.06 ug/L per islet equivalent, p = 0.066) (Fig 4D-E). Expression of Nb-Ft-2-TRPV1^Ca2+^ in islets did not alter insulin content (Fig S3C) or glucose-stimulated insulin release from islets without exposure to a magnetic field (Fig S3 D, E). There was also no effect of the magnetic field on insulin release from beta cells expressing the control mCherry construct (Fig S3F-G). These data confirmed that islets with beta cell expression of Nb-Ft-2-TRPV1^Ca2+^ show increased glucose-stimulated insulin secretion in a magnetic field *ex vivo*.

We next tested whether the Nb-Ft-2-TRPV1^Ca2+^ construct could remotely modulate insulin release *in vivo* leading to changes in blood glucose. In these studies, we delivered the magnetic field (300mT) using a 7 T MRI machine (Fig 4F). Since magnetic field exposure *ex vivo* showed enhanced glucose-stimulated insulin release with a reduced effect on basal insulin release, we reasoned that magnetogenetic activation of beta cells *in vivo* would improve glucose tolerance without effects on basal blood glucose. Consistent with this, baseline blood glucose was similar in Ins1-cre mice that received injections of the DIO-Nb-Ft-2-TRPV1^Ca2+^ or DIO-mCherry AAV in the presence of a magnetic field (Fig 4G). However, in Ins1-cre mice with beta cell expression of Nb-Ft-2-TRPV1^Ca2+^, exposure to the magnetic field during a glucose tolerance test (GTT) resulted in significantly reduced peak blood glucose levels at 15 minutes post-glucose injection (p < 0.01) (Fig 4H), with a significant decrease in the AUC during the first 45 minutes (p < 0.01) (Fig 4I) and for the entire 90 minutes of the GTT (p < 0.05) (Fig 4J) compared to mice with beta cell expression of mCherry. The improvement in glucose tolerance was accompanied by a significant increase in glucose-stimulated insulin secretion 10 minutes after glucose injection, without changes in glucagon release (plasma insulin 10min: 1.41 ± 0.14 µ/L for Ins-cre/Nb-Ft-2-TRPV1^Ca2+^, 1.01 ± 0.11 ug/L for Ins-cre/mCherry, p < 0.05) (Fig 4J-K). There was no effect of the construct on glucose tolerance in animals that were not placed in the MRI machine (Fig S3H). These data show that magnetic activation of pancreatic beta cells after delivery of an AAV expressing Nb-Ft-2-TRPV1^Ca2+^ can increase glucose-stimulated insulin release and improve glucose tolerance *in vivo*.

### Benchtop device for magnetic field delivery regulates insulin release *in vivo*

While an MRI machine can deliver a sufficiently strong field for magnetogenetic activation, this instrument is costly and access can be limited. Thus, a tunable benchtop device capable of delivering a sufficiently strong magnetic field for cell modulation *in vivo* would provide a more cost-effective and simpler alternative to using an MRI. Toward this end, we designed a novel system using paired, water-cooled electromagnets (Helmholtz coils) to generate a relatively uniform magnetic field (Fig 5A-B). The magnetic field was modeled using Matlab (MathWorks) and predicted to have a field strength of 250mT when a current of 496 A was supplied (Fig 5B). In line with these calculations, the strength of the magnetic field at a current of 496 A was measured as 248mT using a Bell 5180 magnetometer and the magnetic field varied linearly with the current. We next tested whether the magnetic field generated by this device could reproduce the effects observed after placement of animals in an MRI. We placed Ins1-cre mice into this device four weeks after injection of DIO-Nb-Ft-2-TRPV1^Ca2+^ AAV or DIO-mCherry control AAV into the pancreas and performed glucose tolerance tests before and after exposure to a magnetic field. We found that animals placed into the device with the field turned on showed significantly improved glucose tolerance to a similar extent as was seen using the MRI (Blood glucose (30 min): 295 ± 42 mg/dL for Nb-Ft-2-TRPV1^Ca2+^, 384 ± 19 mg/dL for mCherry, p < 0.01. Blood glucose (45 min): 233 ± 31 mg/dL for Nb-Ft-2-TRPV1^Ca2+^, 299 ± 17 mg/dL for mCherry, p < 0.05. Mixed effects model, p < 0.03) (Fig 5C-E) and enhanced glucose-stimulated insulin release *in vivo* (Plasma insulin (10min): 1.17 ± 0.13 µg/L for Nb-Ft-2-TRPV1^Ca2+^, 0.82 ± 0.09 µg/L for mCherry, p < 0.05) without effects on plasma glucagon (Fig 5F-G). The GTTs were unchanged when animals were placed inside the device with the current turned off (Fig S3H). These findings are the same as those seen after placing animals in the MRI using the same protocol for Nb-Ft-2-TRPV1^Ca2+^ expression in beta cells. These data validate the utility of this benchtop device for activating cells expressing Nb-Ft-2-TRPV1^Ca2+^ *in vivo*.

**Figure 5:**
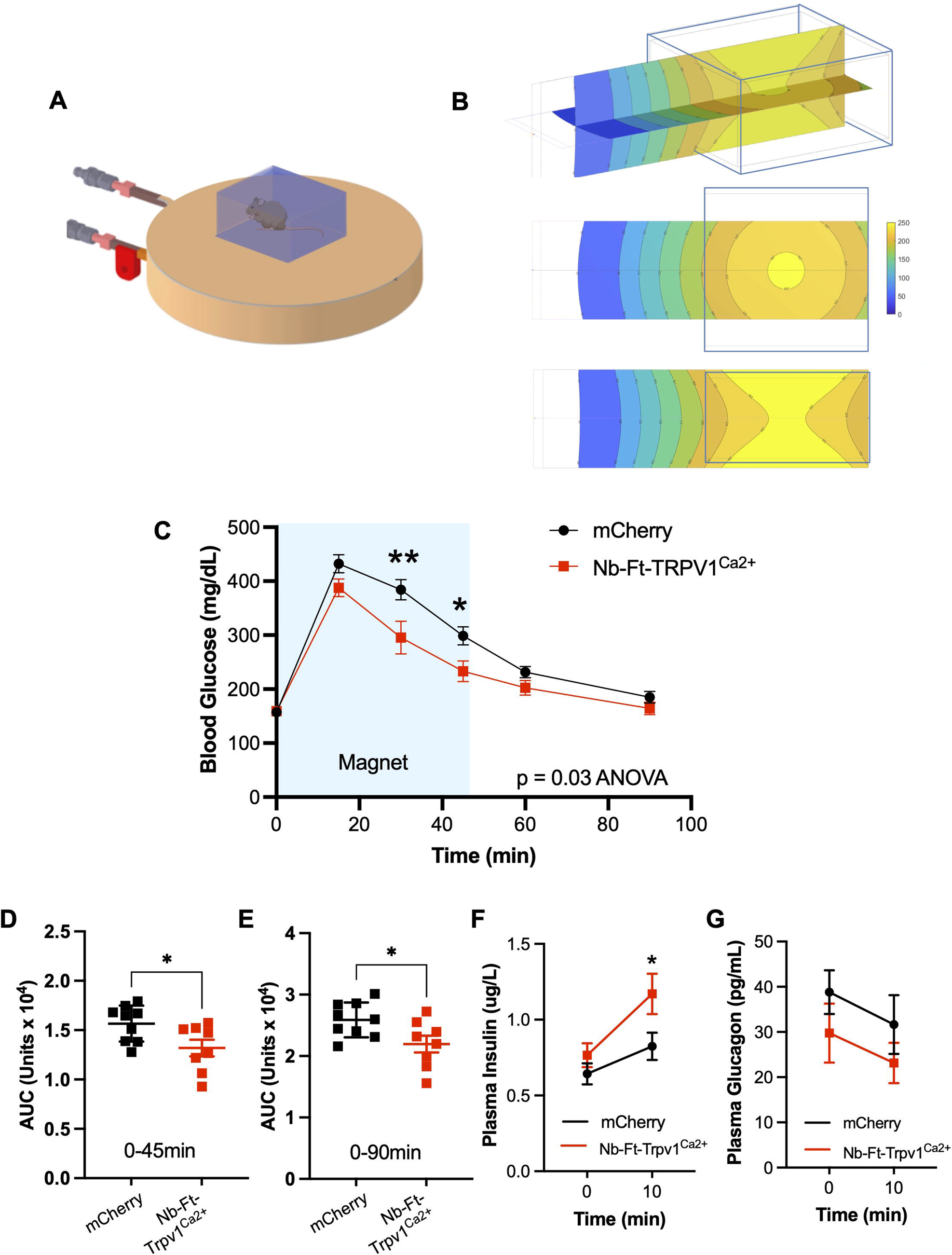
*In vivo* stimulation of Ins1-cre mice with benchtop magnet device. A. Schema of benchtop Helmholtz coil for *in vivo* studies (with upper coil removed for clarity) and *in vivo* chamber. B. Magnetic field in oblique (upper), x (upper) and y (lower) axes with 496A current supplied to the paired coils with position of *in vivo* chamber. Field modeled in Matlab. C. Effects of benchtop magnet treatment (peak 248mT) on blood glucose in Ins1-cre mice with intrapancreatic injection of AAV8-JeT-DIO-Nb-Ft-2-TRPV1^Ca2+^ or AAV8-JeT-DIO-mCherry. Data were analyzed by mixed effects analysis with Sidak’s multiple comparison test. * p = 0.034 Nb-Ft-2-TRPV1^Ca2+^ vs. mCherry, ** p = 0.001 at 30 mins, Nb-Ft-2-TRPV1^Ca2+^ vs. mCherry, * p = 0.02 at 45 mins, Nb-Ft-2-TRPV1^Ca2+^ vs. mCherry. D. Effects of benchtop magnet treatment on cumulative blood glucose during 0-45 mins of glucose tolerance testing (2g/kg) in Ins1-cre mice with intrapancreatic injection of AAV8-JeT-DIO-Nb-Ft-2-TRPV1^Ca2+^ or AAV8-JeT-DIO-mCherry. Data were analyzed by unpaired two-tailed t-test. * p = 0.028 Nb-Ft-2-TRPV1^Ca2+^ vs. mCherry. E. Effects of benchtop magnet treatment on cumulative blood glucose during 0-90 mins of glucose tolerance testing (2g/kg) in Ins1-cre mice with intrapancreatic injection of AAV8-JeT-DIO-Nb-Ft-2-TRPV1^Ca2+^ or AAV8-JeT-DIO-mCherry. Data were analyzed by unpaired two-tailed t-test. * p = 0.029 Nb-Ft-2-TRPV1^Ca2+^ vs. mCherry. F. Effects of benchtop magnet treatment on plasma insulin at 0 and 10 mins during a glucose tolerance test (2g/kg) in Ins1-cre mice with intrapancreatic injection of AAV8-JeT-DIO-Nb-Ft-2-TRPV1^Ca2+^ or AAV8-JeT-DIO-mCherry. Data were analyzed by two-way repeated measures ANOVA with Bonferroni’s multiple comparison test * p = 0.02 Nb-Ft-2-TRPV1^Ca2+^ vs. mCherry at 10 mins. G. Effects of magnet treatment on plasma glucagon at 0 and 10 mins during a glucose tolerance test (2g/kg) in Ins1-cre mice with intrapancreatic injection of AAV8-JeT-DIO-Nb-Ft-2-TRPV1^Ca2+^ or AAV8-JeT-DIO-mCherry. Data were analyzed by mixed measure analysis with Bonferroni’s multiple comparison test p = 0.4 Nb-Ft-2-TRPV1^Ca2+^ vs. mCherry. Data are shown as mean ± SEM. AAV8-JeT-DIO-Nb-Ft-2-TRPV1^Ca2+^ n = 8, AAV8-JeT-DIO-mCherry n = 9.

## Discussion

Systems to transduce magnetic fields for cell activation have been developed by tethering multimodal TRPV channels to ferritin(*8, 9, 15*). In previous studies from our laboratories, we modulated cell activity *in vitro* and *in vivo* using an anti-GFP nanobody to tether TRPV1 to GFP-tagged chimeric ferritin(*9, 16, 20*). We have shown that this approach can be used to modulate calcium-dependent SEAP release from HEK-293T cells(*20*), calcium-dependent insulin expression and release from engineered mesenchymal stem cells and hepatocytes *in vivo*(*16*), and can modulate activity of glucose-sensing neurons *ex vivo* and *in vivo*(*9*). However, this two-component system exceeds the packaging capacity of the most commonly used viral vector, AAV. In order to minimize the size of magnetogenetic actuators, we generated a novel ferritin-binding nanobody to tether TRPV1 to endogenous ferritin. This 2.9 kb construct composed of a TRPV1-anti-ferritin fusion protein can be efficiently packaged into a single AAV that was capable of enhancing glucose-stimulated insulin secretion *in vitro* and *in vivo*. The *in vivo* studies employed a novel device that can deliver a magnetic field that is sufficiently strong for channel gating and is cheaper, and thus more accessible, than an MRI machine used previously in our *in vivo* studies. The development of a system that can be packaged into an AAV and a suitable device for magnetogenetic activation broadens the applicability of the method in animals and potentially clinical settings.

These findings are consistent with many previous studies that have shown that TRP channels tethered to ferritin are gated in the presence of a magnetic field. Published studies have demonstrated magnet treatment of cells expressing ferritin-tethered TRPV1 or TRPV4 can increase intracellular calcium(*10–12, 20, 24*), modulate cell migration(*14*), regulate calcium-dependent gene expression(*16, 20*), modify activity in neural crest cells *in vivo*(*8*), and regulate neural activity to modify reward behavior(*15*) and metabolic regulation(*9*). Our finding that TRPV1 can be tethered to endogenous ferritin using a nanobody and this interaction allows gating of the channel by magnetic field stimulation is also consistent with a prior report showing that endogenous ferritin tethered to TRPV4 by the ferritin-binding domain 5 of kininogen-1 can modulate cell activity(*8, 11, 12*).

Several groups have published empiric data demonstrating a cellular response to magnet treatment of cells expressing TRP channels tethered to ferritin(*8–15, 20*). A number of biophysical and biochemical mechanisms have been proposed for the magnetic activation of multimodal TRP channels tethered to ferritin. The concept that thermal or mechanical forces gated the activation of TRP channels by ferritin was initially questioned on theoretical grounds(*25*). However, subsequent detailed theoretical analyses of physical parameters of ferritin-iron nanoparticles and their interactions with multimodal TRPV ion channels suggest the possibility that alternative thermal or mechanical forces result in channel opening (*13, 24, 26*). A thermal mechanism was further supported by *in vitro* studies demonstrating reduced channel activation with temperature-insensitive TRPV4 channel mutants(*13*). Recent studies from our group and others support the role of an indirect biochemical mechanism for magnetic activation of TRPV1 and TRPV4 channels by ferritin(*11, 12, 20*). Interactions between magnetic fields and ferritin result in local increases in cytosolic free iron that, in turn, generate reactive oxygen species (ROS) to activate ferritin-tagged TRPV1 or V4 channels. This proposed mechanism is supported by *in vitro* imaging studies demonstrating highly localized increases in cytosolic iron and reactive oxygen species, magnetic activation of ROS-sensitive, cold activated TRPA1 channels tethered to ferritin and loss of activation with ROS inhibitors or TRPV1 mutants insensitive to reactive oxygen species(*11, 20, 27*). These mechanisms are not mutually exclusive and both chemical and physical means may contribute to channel opening. While there is abundant evidence from numerous groups confirming the phenomenon, further studies will be needed. Our short, unimolecular TRPV1-ferritin nanobody fusion construct may provide an additional tool for these mechanistic studies. For example, our single construct may reduce the variability associated with co-expressing two constructs (GFP-tagged ferritin and anti-GFP TRPV1), may enabling assessment of mutant TRP channels with defined properties through fusion with the anti-ferritin nanobody or may facilitate the assessment of mechanisms across cells or tissues from multiple species.

Nanobodies offer the advantage of small size, high affinity and high stability across a wide range of conditions(*18*). The ferritin nanobody generated here bound human ferritin with relatively high affinity *in vitro* albeit with lower affinity than for anti-GFP nanobodies. Despite the lower affinity, the anti-ferritin and anti-GFP nanobodies showed equivalent cell activation *in vitro* and *in vivo* in all experiments that were performed. This suggests that the level of expression of the nanobody and ferritin is sufficiently high to lead to complex formation between ferritin and TRPV1. The custom-made ferritin nanobody that was selected bound to both mouse and human ferritin so it could be used effectively in both species. The amino acid sequence of ferritin heavy chain is slightly more conserved through evolution than ferritin light chain, with 93%(*28*) and approximately 85%(*29*) homology between human and mouse ferritin heavy and light chains, respectively. In keeping with this, the mass spectrometry data demonstrated mouse ferritin heavy chain was the major component of the immunoprecipitated nanobody-ferritin complex. The relatively high homology in ferritin heavy chains across many species suggests the anti-ferritin nanobody may effectively bind ferritin of other species. If confirmed, this would expand the ability of the construct to regulate cell activity in other organisms.

Our studies confirmed cell membrane expression and channel function of Nb-Ft-tagged TRPV1^Ca2+^ *in vitro*. As mentioned, despite the lower binding affinity between Nb-Ft and ferritin compared to that between Nb-GFP and GFP, magnet treatment was as effective in activating cells expressing Nb-Ft-TRPV1^Ca2+^/GFP-ferritin as in cells expressing Nb-GFP-TRPV1^Ca2+^/GFP-ferritin. The efficacy of the Nb-Ft-TRPV1^Ca2+^ was confirmed by multiple methods: calcium imaging and calcium-dependent reporter release *in vitro* as well as modulation of endogenous insulin release from pancreatic beta cells *ex vivo* and *in vivo*. AAV delivery of Nb-Ft-TRPV1^Ca2+^ constructs enables *in vivo*, remote, targeted modulation of cell types dispersed across organs, such as beta cells in pancreatic islets. The AAV-compatible magnetogenetic construct combined with advances in AAV serotypes, cell-type specific promoters, splicing and intersectional viral approaches(*30–32*) will greatly expand the questions that can be addressed by magnetogenetics.

Previous studies have demonstrated that effective magnet activation of cells requires close association between TRPV receptors, ferritin and membrane lipids(*11, 16, 20*). Magnet-activated constructs use multimodal transient receptor potential ion channels (TRPV1 or TRPV4) directly or indirectly bound to the iron storage protein, ferritin or a synthetic iron nanoparticle. While the precise mechanisms by which magnetic fields gate the channel is under investigation, several lines of evidence suggest that the channel needs to be in proximity to the magnetically susceptible particle. Recent work has suggested biochemical mechanisms for magnet activation of TRPV channels, relying on interactions between endogenous or chimeric ferritin and oscillating magnetic fields to generate local reactive oxygen species and oxidized lipids that in turn activate TRPV channels(*11, 20*). In this case, rapid diffusion requires that the two moieties be in proximity. Similarly, theoretical calculations suggesting that heat and/or mechanical forces may also increase the probability of channel opening also suggest the need for close proximity between TRP channels and an iron oxide particle(*33*). The use of a small nanobody to link endogenous ferritin to the channel satisfies this requirement for proximity. Consistent with a chemical mechanism for channel activation, recent work in Neuro2A cells expressing TRPV4 tethered to ferritin show similar responsiveness to magnetic field stimulation (180 MHz, 1.6 μT) at 22°, 32° and 37°C with faster activation and decay kinetics at 37°C(*12*).

A range of devices have been used for generating a sufficiently strong magnetic field to gate magnetically-activated constructs. To date, these have largely been designed to address a specific experimental question rather than provide an instrument(s) that can be applied to a broad range of *in vitro* and *in vivo* studies. In this report we developed two additional devices for applying a magnetic field, one using permanent magnets for *in vitro* studies and another using an electromagnet for *in vivo* studies. For longer-term treatment *in vitro* studies, we developed an oscillating permanent magnet array that can be moved closer to or farther from the samples to deliver to cells a customizable, intermittent magnetic field to cells. Permanent magnets, unlike electromagnets, do not disrupt the temperature control in a cell culture incubator, and therefore, allow magnet treatment of cells that require a precise environment, such as islets. Using this approach, we confirmed that beta cell expression of Nb-Ft-TRPV1^Ca2+^ enhanced glucose-stimulated insulin release from mouse islets. We also designed a bench top Helmholtz coil system for *in vivo* studies. Unlike magnetic fields from MRI coils or arrays of powerful permanent magnets used for previous *in vivo* studies, this system can deliver continuous or pulsed fields with controlled field strength. The current system does not require expensive cooling equipment so it provides a relatively cost-effective method for *in vivo* studies in mice but could be modified with larger coils and cooling systems to treat other species.

This bench top device was as effective as the field outside an MRI coil at regulating beta cells expressing Nb-Ft-TRPV1^Ca2+^ to increase plasma insulin and improve glucose tolerance. While a magnet field was sufficient to potentiate glucose-stimulated insulin release ex vivo and improve glucose tolerance *in vivo*, magnetically induced insulin release from beta cells at lower glucose levels was not significantly increased. However, we found only relatively low levels of expression of the construct after injection of AAV8 and hypothesize that *in vivo*, this relatively low expression level was not sufficient to alter blood glucose with magnet treatment in the absence of a glucose challenge. Consistent with this, gap junction coupling is reported to attenuate beta cell activity under basal conditions by preventing individually depolarized beta cells from reaching the threshold for insulin secretion, but enhances beta cell activity in an islet as glucose levels rise(*34, 35*). With the use of novel AAV serotypes(*36*) it may be possible to improve beta cell construct expression and so lower both basal and glucose-stimulated plasma glucose with magnet treatment.

Our AAV-compatible magnetogenetic construct and a low-cost device for magnetogenetic activation expands the applicability of the method for both preclinical studies and, potentially, translational use. In parallel studies performed using the approaches developed in this paper, we have shown that AAV delivery of activating and inhibitory versions of the Nb-Ft-TRPV1 construct can modulate neural activity *in vivo* to replicate or attenuate motor defects in mouse models of Parkinson’s disease(*37*). The combination of novel AAV serotypes for targeted delivery of magnetogenetic constructs and the use of novel devices for generating magnetic fields could offer rapid, more specific control of neural populations than existing neuromodulation devices. This could be advantageous for translational uses such as pain control(*38, 39*), neuromodulation for incontinence(*40, 41*), or as an alternative to vagal nerve stimulation for epilepsy(*42*) or treatment-resistant depression(*43*). In addition, this approach could take advantage of the growing number of devices already approved by the FDA to deliver magnetic fields to the central nervous system such as transcranial magnetic stimulators(*44*) and peripheral nerve stimulators(*45*). Since expression of Nb-Ft-TRPV1 sensitizes cells to magnetic fields, magnetogenetic constructs also offer the potential to expand the range of disorders amenable to transcranial magnetic stimulation treatment by minimizing the field strength necessary to elicit an effect.

In summary, we have generated a new single component system for magnetic activation of mouse and human cells suitable for expression in an AAV together with a series of instruments for generating a magnetic field of sufficient strength *in vitro* and *in vivo*. We have applied these tools to remotely regulate cell activity *in vitro* and also modulate the function of pancreatic beta cells, a distributed endocrine population, *in vivo* to improve glucose control in mice. These tools offer a platform for remote cell modulation in cultured cells for *in vivo* physiology, in human and mouse as well as potentially other species. These approaches are complementary to optogenetics and chemogenetics and can be used to dissect the physiological roles of defined cell populations.

## MATERIALS AND METHODS

### Ferritin nanobody generation

#### Immunization of a llama

A llama was injected subcutaneously with human spleen ferritin (250 μg) (Lee Biosolutions, MO) on days 0, 7, 14, 21, 28 and 35. Gerbu LQ#3000 was used as adjuvant. On day 40, anticoagulated blood was collected for lymphocyte preparation.

#### Construction of a VHH library

A VHH library was constructed and screened for the presence of antigen-specific nanobodies. To this end, total RNA from peripheral blood lymphocytes was used as template for first strand cDNA synthesis with oligo(dT) primer. Using this cDNA, the VHH encoding sequences were amplified by PCR, digested with PstI and NotI, and cloned into the PstI and NotI sites of the phagemid vector pMECS. A VHH library of about 10^8^ independent transformants was obtained. About 87% of transformants harbored the vector with the right insert size.

#### Isolation of antigen-specific nanobodies

The library was subject to three consecutive rounds of panning on solid-phase coated human ferritin (200 µg/ml, 20 μg/well). The enrichment for human ferritin-specific phages was assessed after each round of panning by comparing the number of phagemid particles eluted from antigen-coated wells with the number of phagemid particles eluted from negative control wells blocked without antigen. These experiments suggested that the phage population was enriched about 20-fold, 20-fold, and 8 × 10^2^-fold for human ferritin-specific phages after 1^st^, 2^nd^ and 3^rd^ rounds of panning, respectively. 190 colonies from 1^st^ & 2^nd^ rounds were randomly selected and analyzed by ELISA for the presence of human ferritin-specific nanobodies. This first round of ELISAs was used to screen nanobodies in crude periplasmic extracts and the relative intensity of the ELISA signal may not reflect the relative quality of the nanobodies. In these experiments the differences in ELISA signals between different nanobodies may be related to variables such as the amount of nanobody, rather than to nanobody quality such as affinity or actual yield. Out of these 190 colonies, 133 colonies scored positive for the presence of nanobodies to human ferritin in this assay (46 & 87 from 1^st^ & 2^nd^ rounds, respectively). The human ferritin used for panning and ELISA screening was the same as the one used for immunization. Based on sequence data, the ELISA-positive colonies represented 59 different nanobodies belong to 22 different CDR3 groups.

Next, the library was subject to three consecutive rounds of panning on solid-phase coated with mouse ferritin purified from mouse liver (200 µg/ml, 20 μg/well). The enrichment for mouse ferritin-specific phages was assessed as above. These experiments suggested that the phage population was enriched about 10-fold, 10-fold and 5 × 10^2^-fold for mouse ferritin-specific phages after 1^st^, 2^nd^ and 3^rd^ rounds of panning, respectively. 190 colonies from 2^nd^ & 3^rd^ rounds were randomly selected and analyzed by ELISA for the presence of mouse ferritin-specific nanobodies in their periplasmic extracts (ELISA using crude periplasmic extracts including soluble nanobodies). Out of these 190 colonies, 54 colonies scored positive in this assay (1 & 53 from 2^nd^ & 3^rd^ rounds, respectively). Based on sequence data, the ELISA-positive colonies represented 19 different nanobodies belonging to 6 different CDR3 groups. Two nanobody CDR3 groups bound both human and mouse ferritin.

### Purification of recombinant nanobody protein

pMECS vectors encoding anti-ferritin nanobodies were transformed into electrocompetent E.coli WK6 cells. Colonies were picked and grown in 10-20 ml of LB + ampicillin (100 µg/ml) + glucose (1%) at 37°C overnight. 1ml of preculture was added to 330 ml TB-medium (2.3 g KH_2_PO_4,_ 16.4 g K_2_HPO_4_.3H_2_O,12 g Tryptone, 24 g Yeast and 4 ml 100% glycerol supplemented with 100 µg/ml Ampicillin, 2mM MgCl_2_ and 0.1% glucose and grow at 37°C. Nanobody expression was induced by addition of IPTG (1mM). The culture was incubated at 28°C with shaking overnight.

The cells were pelleted for 8 minutes at 8000 rpm and resuspended in 12 ml TES (0.2 M Tris pH8.0, 0.5 mM EDTA, 0.5 M sucrose) before shaking for 1 hour on ice. A further 18ml of TES (diluted 4 fold in water) was added and incubated for a further hour on ice with shaking then centrifuged for 30 mins at 8000rpm, at 4°C. The supernatant containing the nanobody proteins was then purified by immobilized metal-ion affinity chromatography (IMAC) using HIS-Select® Cobalt Affinity Gel (Sigma H8162) according to the manufacturer’s instructions. The amount of protein was estimated at this point by OD_280_ measurement of eluted sample. 14 nanobodies were successfully purified.

### Characterization of nanobody affinity to human ferritin by ELISA

ELISA plates were coated with 1µg/mL human spleen ferritin (Lee Biosolutions, MO) in PBS overnight at 4°C. Plates were then washed 5 times with washing buffer (PBS+0.05% Tween-20), followed by blocking with PBS+ 1% BSA at room temperature for 2 hrs. Serial dilutions of nanobodies or BSA (100ul) were added to the ELISA plates and incubated for 2 hours at room temperature then washed 5 times with buffer. Anti-HA-HRP antibody (100ul, Miltenyi biotech, 130-091-972, 1:1000 dilution) was added to each well and incubated for 1 hr at RT before a further 5 washes. TMB substrate (100ul) was added to each well and the reaction stopped by addition of 100 µL 2M H_2_SO_4_ after 10 mins. OD450 was measured using a microplate reader (BMG Labtech CLARIOstar plate reader).

### Characterization of anti-ferritin nanobodies by immunoprecipitation

Nanobody sequences were amplified by PCR with the specific primers (NbFt-F: gcaagatctgccaccatggcc CAGGTGCAGCTGCAGGAG; NbFt-R:gcaaagcttggatccAGCGTAATCTGGAACATCGTATGGGTAtgcggccgctgagga) and subcloned into the retroviral vector pMSCV (Clontech, Takara Bio, US). HEK-293T cells were grown in 10cm plates and cotransfected with plasmids expressing GFP-mFerritin and anti-ferritin nanobody clones using PEI (DNA:PEI=1:3). Cells were harvested after 48 hours and lysed with 500 µl of lysis buffer (50 mM Tris, pH 7.5, 300 mM NaCl, 1 mM EGTA, 1 mM EDTA, 1% NP-40, 0.1% SDS) and protease inhibitor (cOmplete EDTA-free, Roche) then incubated for 15 mins with agitation at 4°C before centrifugation at 14,000 rpm for 20 mins. The supernatant was transferred to a new tube with 30 µl HA.11 antibody-conjugated agarose beads (Biolegend, 900801) added to each sample. The beads and supernatant were incubated at 4°C overnight with rotation and then spun at 3000rpm for 2 mins. Beads were then washed 3 times with lysis buffer before proteins were eluted with 20 µl of 4x Laemmili buffer and heated at 95°C for 5 minutes. Eluted proteins were examined by SDS-PAGE.

Samples were loaded onto a 10 well SDS-PAGE gel and run for 10-15mm then fixed in 46% methanol/7% glacial acetic acid for 1 hour. The gel was then stained for 1 hour in 0.1% Coomassie blue R-250 in 46% methanol/ 7% glacial acetic acid before destaining in ultrapure water. Bands were then excised and analyzed by mass spectrometry.

### Mass spectrometry

Proteins were reduced (10 mM DTT, EMD Millipore) and alkylated (30 mM iodoacetamide, Sigma), followed by digestion with Endoproteinase LysC (Wako Chemicals) and trypsin (Promega). Reaction was halted by addition of trifluroacetic acid and peptides were solid phase extracted (*46*) and analyzed by reversed phase nano-LC-MS/MS (Dionex 3000 coupled to a Q-Excative Plus, Thermo Scientific). Data were quantified and searched against a Uniprot human database concatenated with a mouse GFP-Ferritin sequence using ProteomeDiscoverer 1.4/Mascot 2.5. Oxidation of methionine and protein N-terminal acetylation were allowed as variable modifications. Cysteines were considered as fully carbamidomethylated. Two missed cleavages were allowed. Peptide matches were filtered using a percolator(*47*) calculated false discovery rate of 1%. The average area of the three most abundant peptides per protein were used as a proxy for protein abundance (*48*). For the highly homologous ferritin heavy proteins, only peptides unique to either the mouse or the human forms were used in access abundance. Peptides specific to the N-terminal GFP moiety of the over expressed mouse ferritin was not used to calculate abundance of mouse ferritin but were recorded separately.

### Isothermal titration calorimetry

We performed an isothermal titration calorimetry (ITC) assay using a MicroCal Auto-iTC200 instrument (Malvern Panalytical). In each experiment, 10 μM human spleen ferritin in the cell was titrated with 75 μM NbFt2 in the syringe at 25 °C. The titration sequences included a single 0.4 μL injection, followed by 19 injections of 2 μL each, with a 150-sec interval between injections, stirring rate of 750 rpm and reference power of 10 μcal/sec. Data were analyzed with Origin analysis software.

### Constructs

The expression vectors for calcium dependent release of insulin and SEAP, pCMV-TRPV1^Ca2+^ were generated as previously described(*20, 21*). To generate Ca^2+^-dependent SEAP reporter, the SEAP coding fragements were obstained by cutting pYSEAP (addgene 37326) with HindIII and HpaI and inserted into pSRE-CRE-NFAT-insulin(*21*) at the HindIII and HpaI sites.

To express Nb-FT in mammalian cells, a PGK promoter-driven mCherry expression cassette and the WPRE element were inserted into pMSCV (Clontech, Takara Bio) at the XhoI and ClaI sites to generate an empty vector (pMSCV-mCherry-WPRE). The anti-ferritin nanobody coding sequences were amplified from phage display plasmids with specific primers: 5’gcaagatctgccaccatggccCAGGTGCAGCTGCAGGAG and 5’gcaaagctt ggatccAGCGTAATCTGGAACATCGTATGGGTAtgcggccgctgagga, digested with BglII and HindIII and ligated into pJFXY21 at the BglII and HindIII sites. The resulting plasmids were named pMSCV-Nb-Ft-2, -9, -10, -14 and -17-TRPV1^Ca2+^. The NbGFP fragment was amplified from MSCV-αGFP-TRPV1-2A-GFPferritin(*16*) with specific primers: 5’gcaagatctgccaccatggcc CAGGTGCAGCTGCAGGAG and 5’ tcagacgtcggccactgcggccgcTGAGGAGACGGTGACCTGGGTC, digested and subcloned into pNb-Ft-2-TRPV1^Ca2+^ at the BglII and NotI sites to generate plasmid pNb-GFP-TRPV1^Ca2+^.

The pMSCV-Nb-GFP-TRPV1^Ca2+^-T2A-GFP-mFerritin plasmid was generated by subcloning the Nb-GFP-TRPV1^Ca2+^-T2A-GFP-mFerritin fragment cut from MSCV-αGFP-TRPV1-2A-GFPferritin(*16*) with NotI and EcoRI into a modified pMSCV-WPRE vector at the NotI and MfeI. NbFT sequences were amplifed from plasmids pMSCV-Nb-Ft-2, -9, -10, -14 and -17-TRPV1^Ca2+^ with specific primers: 5’ gtgatgcatGCCACCATGGCCCAGGTGCAG and 5’ gatgctagcgccAGCGTAATCTGGAACATCG, digested with NsiI and Nhel and inserted into pJFXY16 at the NsiI and AvrII sites. The resulting constructs were named pMSCV-Nb-Ft-2, -9, -10, -14 and -17-TRPV1^Ca2+^-T2A-GFP-mFerritin.

The pMSCV-GFP-mFerritin vector was created by digesting pMSCV-Nb-GFP-TRPV1^Ca2+^-T2A-GFP-mFerritin with BglII to remove the NbGFP-TRPV1-T2A fragment and self-ligated.

To make pAAV-hSyn-Nb-FT-2-TRPV1^Ca2+^, the 2981-bp PCR product containing Nb-FT-2-TRPV1^Ca2+^was amplified from pMSCV-Nb-Ft-2-TRPV1^Ca2+^-T2A-GFP-mFerritin. using primers 5’AGCGCAGTCGAGAAGGTACCGGATCCCCCGGTCGCCACCACTAGTATGGCCCAGGTGCAGCTGC and 5’-TTATCGATAAGCTTGATATCGAATTCTTACTTCTCCCCTGGGACCATG, digesting with BamHI and EcoRI and cloning into BamHI/EcoRI-digested pAAV-hSyn-mCherry (Jimenez-Gonzalez et al, 2021, NBME).

To make pAAV-hSyn-RCaMP, a 1441bp PCR product containing jRGECO1a, a red fluorescent calcium sensor protein (RCaMP) was amplified from pAAV.Syn.Flex.NES-jRGECO1a.WPRE.SV40 (Addgene cat#100853) using primers 5’-ACTAGTATGCTGCAGAACGAGCTTGC and ACCGGTCTACTAGTCTCAATTGTCACTTCGCTGTCATCATTTGT, digested with SpeI, and cloned into pAAV-hSyn-DIO-MCS in the forward direction so that RCaMP could be expressed in the absence of Cre recombinase. pAAV-hSyn-DIO-MCS is pAAV-hSyn-DIO-hM3D(Gq)-mCherry (Addgene cat# 44361) which had previously had the BsrGI/NheI region containing hM3D(Gq)-mCherry replaced with the following multiple cloning site: TGTACAACTAGTACCGGTTCGCGAGCATGCCCTAGGGCTAGC.

To make pAAV-Jet-DIO-Nb-FT-2-TRPV1^Ca2+^, a 242-bp PCR product containing the JeT promoter (Tornoe et al. Gene 297 (2002) 21–32) was amplified from pAAV-HI-Jet-mCherry (a gift of Michael Kaplitt, Weill Cornell) using primers 5’-CTGCGGCCGCACGCGTGTACCATTGACGAATTCGGGCG and 5’-ACTCTAGAGGATCCGGTACCTGTCAAGTGACGATCACAGGG and cloned into MluI/KpnI digested pAAV-hSyn-DIO-MCS (see RCaMP expression vector above) to make pAAV-JeT-DIO-MCS. Next, the 2943bp SpeI/EcoRV fragment of pAAV-hSyn-Nb-FT-2-TRPV1^Ca2+^ containing Nb-FT-2-TRPV1^Ca2+^ was ligated to NruI/AvrII digested pAAV-JeT-DIO-MCS in a cre-dependent orientation. To make pAAV-JeT-DIO-mCherry, 741-bp PCR product amplified from pCMV-mCherry-C1 (Clontech Laboratories, Inc.) using primers 5’-ATTACCGGTGTTAACATGGTGAGCAAGGGCGAGGA and 5’-TAAACCGGTCTTAAGTTACTTGTACAGCTCGTCCATG was cloned into AgeI-digested pAAV-JeT-DIO-MCS.

### Adeno-Associated Virus Preparation

pAAV-Jet-DIO-Nb-FT-2-TRPV1^Ca2+^ and pAAV-Jet-DIO-mCherry were packaged into adeno-associated viral serotype 2.8 by Virovek (Hayward, CA).

### Cell culture and *in vitro* studies

Human embryonic kidney cells (HEK-293T, (ATCC^®^ CRL-3216^™^) mycoplasma testing and STR profiling performed by ATCC) were cultured in Dulbecco’s modified eagle medium with 10% fetal bovine serum (Gibco, Carlsbad, CA) at 37°C and 5% CO_2_. Neuro 2A cells (ATCC CCL-131, mycoplasma testing and STR profiling performed by ATCC) were grown in Eagles’ minimum essential medium with 10% fetal bovine serum (Gibco) at 37°C and 5% CO_2_.

#### Immunocytochemistry studies

HEK-293T or Neuro 2A cells were cultured on 12-mm cover glass (Fisher Scientific, Pittsburgh, PA) coated with fibronectin (Sigma Aldrich, cat#F1141). Cells were transfected with constructs expressing pAAV-hSyn-Nb-FT-2-TRPV1^Ca2+^ or pCMV-TRPV1^Ca2+^ 24 h after plating using X-tremeGENE^™^ 9 DNA Transfection Reagent (Millipore Sigma) according to manufacturer’s specifications. Cells were stained 48-72hrs after transfection.

#### Magnet treatment studies using calcium-dependent reporters

HEK-293T cells were cultured on 12-mm cover glass and transfected with pMSCV-Nb-Ft-TRPV1^Ca2+^ or pMSCV-Nb-GFP-TRPV1 and GFP-mFerritin and either calcium-dependent SEAP construct(*20*) or calcium-dependent insulin construct(*21*). Holotransferrin (2 mg/ml, Sigma) was added to cells 24 hours after transfection. 24 h prior to the study, cells were placed in 1% FBS medium at 32°C to ensure minimal activation of TRPV1 and calcium dependent pathways. Cells were incubated in 300 µl of calcium imaging buffer at room temperature (control) or in an oscillating magnetic field (465kHz, 30mT) at room temperature. After 60 mins, the supernatant was removed, spun to remove cells and assayed for secreted alkaline phosphatase or insulin. Screening studies with SEAP production were repeated twice with 4 replicates. Validation studies with calcium dependent insulin production were repeated at least 3 times with at least 3 replicates.

#### Calcium imaging studies

Neuro2A cells were cultured on cover glass and transfected with pAAV-hSyn-Nb-FT-2-TRPV1^Ca2+^ with or without pMSCV-GFP-mFerritin as described above with holotransferrin added 24 hours after transfection. Cells were placed at 32°C 24 hours before testing. Cells were loaded with Fluo-4 3µM (Invitrogen) in the presence of sulfinpyrazone 500 µM (Sigma) for 45-60 min at 32°C then washed and incubated for 15-30 min in sulfinpyrazone in PBS. HEK-293T cells were cultured on cover glass and transfected with pAAV-hSyn-RCaMP alone or combined with pAAV-hSyn-Nb-FT-2-TRPV1^Ca2+^ or pCMV-TRPV1^Ca2+^. Cells were then placed in glass bottom dishes with calcium imaging buffer at 29-31°C for imaging. Imaging was performed using a Deltavision personal DV imaging system (Applied Precision, Inc., Issawaq, WA) equipped with a custom-made ceramic lens and softWoRx imaging station. All other calcium imaging experiments were performed on an inverted Zeiss Axio Observer Z1 microscope (Carl Zeiss, Oberkochen GER). Cells were imaged before and during RF treatment, before and during application of a permanent magnet, before and after treatment with capsaicin (1 μM, Sigma). Imaging was performed on at least three occasions for each condition. Image analysis was performed using Image J. Briefly regions of interests (ROI) were selected for background or cells and mean intensity was calculated for every ROI in each image. Calcium responses were quantified as fluorescence intensity normalized to baseline fluorescence.

### Immunocytochemistry

Immunocytochemistry (ICC) and immunohistochemistry (IHC) were used to confirm cell surface expression of TRPV1. Live cells were incubated with rabbit anti-TRPV1 (extracellular) polyclonal antibody (Thermo Fisher Scientific cat#PA5-77361) (1:50) in culture medium for 10 min at 37 °C followed by 5 washes with RT culture medium. Cells were then incubated with goat anti-rabbit Alexa Fluor 568 or 633 (1:1000,) for 10 min at 32 °C followed by a further 5 washes with RT culture medium. Cells were then fixed in 3.7% paraformaldehyde in Hank’s balanced salt solution (HBSS) for 30 min at room temperature and washed 5 times in HBSS before mounting using Fluoromount with DAPI (Southern Biotech, Birmingham, AL). Images were acquired using a Zeiss LSM 880 inverted confocal microscope (Carl Zeiss, Oberkochen GER).

### Electrophysiology recordings of N2A cells expressing pAAV-hSyn-Nb-FT-2-TRPV1^Ca2+^ construct

Neuro2A cells were cultured on uncoated petri dishes and were transfected with pAAV-hSyn-Nb-FT-2-TRPV1^Ca2+^ (as above). Electrophysiological recordings were performed 24-48 hours after transfection in a bath solution consisting of 10mM Hepes p7.4, 140 mM NaCl, 5 mM EGTA and were imaged using a Nikon eclipse DIC microscope at x20 magnification. Pipettes of borosilicate glass (Sutter Instruments; BF150-86-10) were pulled to ∼3-5 MW resistance with a micropipette puller (Sutter Instruments; P-97) and polished with a microforge (Narishige; MF-83). The pipette was filled with identical bath solution. Recordings were obtained with an Axopatch 200B amplifier (Molecular Devices), filtered at 1 kHz and digitized at 10 kHz (Digidata 1440; Molecular devices). Recordings were made in the outside-out excised patch configuration after gigaseals were obtained and currents were recorded by voltage ramp protocols from −100 to +100mV with and without 10 mM capsaicin addition by perfusion (ALA Scientific VM8 manifold perfusion system).

### Devices

#### Oscillating permanent magnet array for cell culture

Magnetic oscillation treatment of cell samples in 96-well plates was conducted using a custom-built oscillator design to fit in standard CO_2_ incubators. The oscillator is composed of a vertically-oriented linear stepper motor (Haydon Kerk Pittman), driven by an IDEA programmable stepper motor controller (Haydon Kerk Pittman), that oscillates a platform with 12 cylindrical, axially magnetized N52 neodymium magnets (6.35 mm diameter by 25.4 mm thick, K&J Magnetics) towards and away from a platform where a 96-well plate with samples is placed, as shown in Figure 4C and S3B. The magnets are arranged such that each is aligned directly below the center of a well with a spacing of one non-treated well between each treated well. During a single oscillation of the magnetic stimulator, the magnets are raised 10 mm from the starting position to be approximately 3.5 mm from the cell layer growing in the well to apply a magnetic stimulation with a peak field strength of 122 mT (Figure S3) before being lowered back down to their starting position, which only has a peak field strength of 15-13 mT (Figure S3). This oscillation was programmed to have a period of 1 s and was repeated continuously for a 30 min treatment window for a total of 1800 magnetic stimulation pulses at 32°C and 5.0% CO_2_. The strength of the magnetic field applied by array of cylindrical magnetics was determined using the Wolfram Mathematica with the Radia software package (European Synchrotron Radiation Facility).

#### Helmholtz coil for in vivo studies

The device consists of 2 copper coils of 65 turns each (5 layers of 13 turns) spaced with 2.5” between them. The coils were wound by Everson Tesla (Nazareth, PA) out of 3/8” square copper tubing with a 7/32” inner diameter for water cooling. The coils are driven by a TSA-10-900 DC power supply (Magna-Power Electronics, Flemington, NJ) controlled by custom control software written in LabVIEW (National Instruments, Austin, Texas). The magnetic field was calibrated using a Bell 5180 magnetometer (FW Bell, Milwaukie, OR). The peak magnetic field was found to be 248 mT when a current of 496A was supplied. As expected, the magnetic field varied linearly with the current and was controllable with 8Hz refresh rate. The magnetic field was modeled using Matlab (MathWorks, Natick, MA).

### Animal studies

Ins1-cre (Jackson Laboratories, Bar Harbor ME; #026801) male mice 8-12 weeks of age were maintained on a 12h/12h light-dark cycle with controlled humidity and temperature, with access to food *ad libitum*. Mice were injected with 5×10^11^ vg of AAV8-JeT-DIO-Nb-FT-2-TRPV1^Ca2+^ or AAV8-JeT-DIO-mCherry via intrapancreatic injection(*49*) and *in vivo* and *ex vivo* studies were performed 4 weeks post-surgery. Animal care and experimental procedures were performed with the approval of the Animal Care and Use Committee of The Rockefeller University and Icahn School of Medicine at Mount Sinai under established guidelines.

#### Islet Isolation and Ex Vivo Studies

Islets from Ins1-cre mice injected with AAV8-JeT-DIO-Nb-FT-2-TRPV1^Ca2+^ or AAV8-JeT-DIO-mCherry were isolated by collagenase digestion at 37°C and histopaque gradient separation as previously described(*50–52*). Per animal, three replicates of 15-20 islet equivalents were placed into cell culture inserts in a 24-well plate. At the same time, another two replicates of 15-20 islet equivalents were placed into cell culture inserts in a separate 24-well plate. Islets were cultured in KREBS buffer + 1% BSA + 5.5mM glucose at 32°C for 1h before initiation of glucose-stimulated insulin secretion (GSIS) experiments. GSIS studies were conducted as previously described(*53*) with the triplicate plate positioned on the magnet stepper and the duplicate plate positioned away from any magnetic field. Briefly, islets were incubated in 5.5mM (low) glucose for 30min followed by incubation in 11 mM (high) glucose for 30 min, +/− magnet treatment. Magnet treatment consisted of the following: magnet oscillation at 1 Hz, generating an oscillating magnetic field with minimum and maximum strengths of 15 mT and 122 mT. After incubation in 11 mM glucose, islets were washed and digested with 0.1N NaOH for measurement of total insulin content. Supernatant was collected after all incubation steps for measurement of insulin secretion. All incubations were performed at 32°C + 5% CO2, using KREBS buffer + 0.1% BSA with varying glucose concentrations. Insulin secretion was measured using a mouse insulin ELISA kit (Mercodia, Uppsala SWE; #10-1249-01).

#### In vivo Metabolic Studies

Ins1-cre mice were injected with AAV8-JeT-DIO-Nb-FT-2-TRPV1^Ca2+^ or AAV8-JeT-DIO-mCherry as described above. Four weeks after surgery, mice were habituated to handling, test chambers (20 cm x 8 cm x 8 cm for MRI treatment, 10 cm x 10 cm x 4 cm opaque chambers) and test room for 3 – 5 days before metabolic studies. The magnetic field for *in vivo* studies was generated by the superconducting electromagnetic MRI field from a BioSpec High-Field 7 Tesla MRI Scanner (Bruker, Cartaret NJ), with a field strength of 300mT, or the custom-made benchtop device. Baseline measurements and intraperitoneal glucose tolerance tests (IPGTT) were administered in the afternoon after a 6h fast. For baseline, blood glucose was measured at times 0, 30 and 60min. For IPGTT, 2 g/kg of D-glucose (MilliporeSigma, St. Louis MO; G8270) in saline was injected at time 0, with blood glucose measured at times 0, 15, 30, 45, 60 and 90min. Glucose was measured on blood collected from the tail using the Contour Next EZ glucose meter (Bayer, Leverkusen GER). Plasma insulin and glucagon were measured during the IPGTT studies at time 0 (immediately preceding glucose injection) and time 10. Plasma levels of insulin and glucagon were determined by ELISA (Mercodia, Insulin: #10-1247-01; Glucagon: #10-1281-01).

#### Tissue Collection and Immunostaining

Mice were perfused with 10% neutral-buffered formalin (Epredia, Kalamazoo MI; #9400-1) and pancreata were harvested. Pancreata were post-fixed in 10% neutral-buffered formalin overnight, immersed in 30% sucrose (MilliporeSigma; #50389) overnight, then embedded in O.C.T. Compound (Thermo Fisher Scientific, Waltham MA; #23-730-572), frozen at −80°C, and sectioned at 10µm thickness. Pancreata were stained overnight for HA-Tag (Cell Signaling, Danvers MA; #3724) at 1:500 dilution + insulin (R&D Systems, Minneapolis MN; #MAB1417) or glucagon (MilliporeSigma; G2654) at 1:1000 dilution. Subsequent secondary antibodies used were Alexa Fluor 647 anti-rabbit (Jackson ImmunoResearch, West Grove PA; #711-605-152), Alexa Fluor 488 anti-rat (Thermo Fisher Scientific; #A21208) and Alexa Fluor 488 anti-mouse (Thermo Fisher Scientific; #21202). Tissues were stained for DAPI (Thermo Fisher Scientific; #D3571) at 1:10,000 dilution and visualized using a Zeiss LSM 880 inverted confocal microscope (Carl Zeiss, Oberkochen GER). Image analysis was performed using the Imaris spots tool as described previously(*54*).

### Statistical Analyses

Data are expressed as means ± SEM. Analyses were performed using GraphPad Prism version 8.3.1. Normality was assessed using the Shapiro-Wilk normality test, and if normality assumption was met (alpha = 0.05), statistics were performed using Student’s unpaired two-tailed t-test for comparison between 2 groups, one-Way Analysis of Variance (ANOVA) with Tukey’s multiple comparison test between multiple groups and two-way repeated measures ANOVA, followed by Sidak’s or Bonferroni’s multiple comparison test for analysis of multiple groups over time. If normality was not met, statistics were performed using Mann-Whitney U test for comparison between 2 groups and Kruskal-Wallis test followed by Dunn’s multiple comparisons test for comparison between multiple groups. P-values <0.05 were considered to be significant.

## Data and Resource Availability

Nanobody protein sequences generated in this study are provided in the manuscript. Key plasmids generated in this study are available from the corresponding authors on request. All data needed to evaluate the conclusions in the paper are present in the paper and/or the Supplementary Materials.

## Acknowledgements

We would like to thank Mladen Barbic, Roberta Marongiu and Michael Kaplitt for helpful discussions. MJG is supported in part by the Naomi Berrie Diabetes Center Russell Berrie Foundation Award. J.M.F acknowledges support from JPB foundation. SAS was supported by the American Diabetes Association Pathway to Stop Diabetes Grant ADA #1-17-ACE-31, and in part by grants from the National Institutes of Health (R01NS097184, OT2OD024912, R01DK124461), Department of Defense (W81XWH-20-1-0345 and Discovery Award # W81XWH-20-1-0156 to MJG). Microscopy and image analysis were performed at the Microscopy Core at the Icahn School of Medicine at Mount Sinai. The authors wish to thank the NIDDK supported Einstein-Sinai Diabetes Research Center (DRC) (P-30 DK020541).

Figures were created with BioRender.com

## Author contributions

LEP, RL, LK, and GV designed studies, performed experiments, analyzed data and contributed to the writing of the manuscript. XY, GH, HM, DG, MB, PW, and MJG performed experiments and reviewed the manuscript. AGO and JD designed studies and contributed to the writing of the manuscript. JMF and SAS designed studies, analyzed data and wrote the manuscript. All authors discussed the results and edited the manuscript.

