## Supplementary figures and images for "Magnetogenetic cell activation using endogenous ferritin"

### Supplemental figures 23.6.20

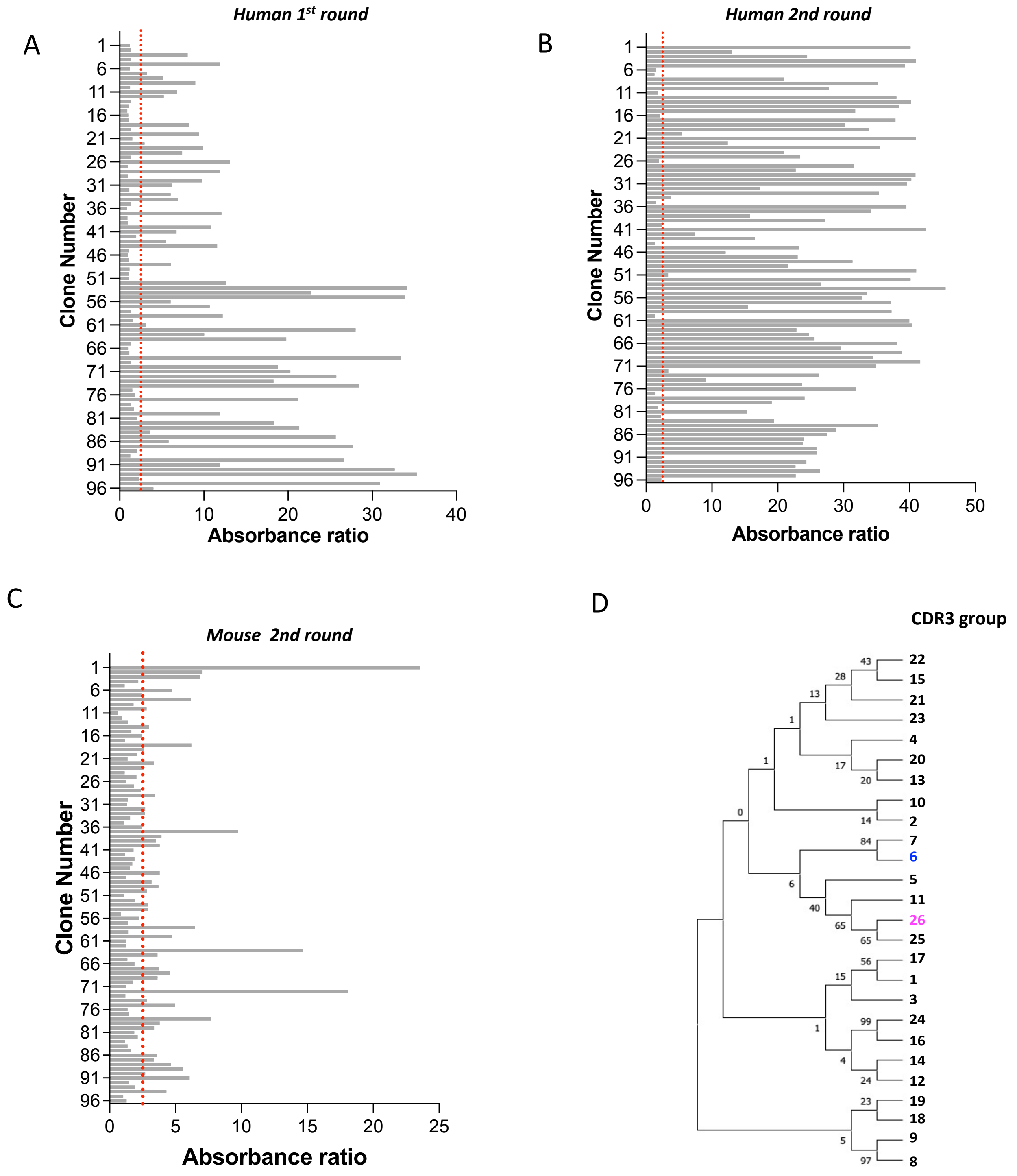

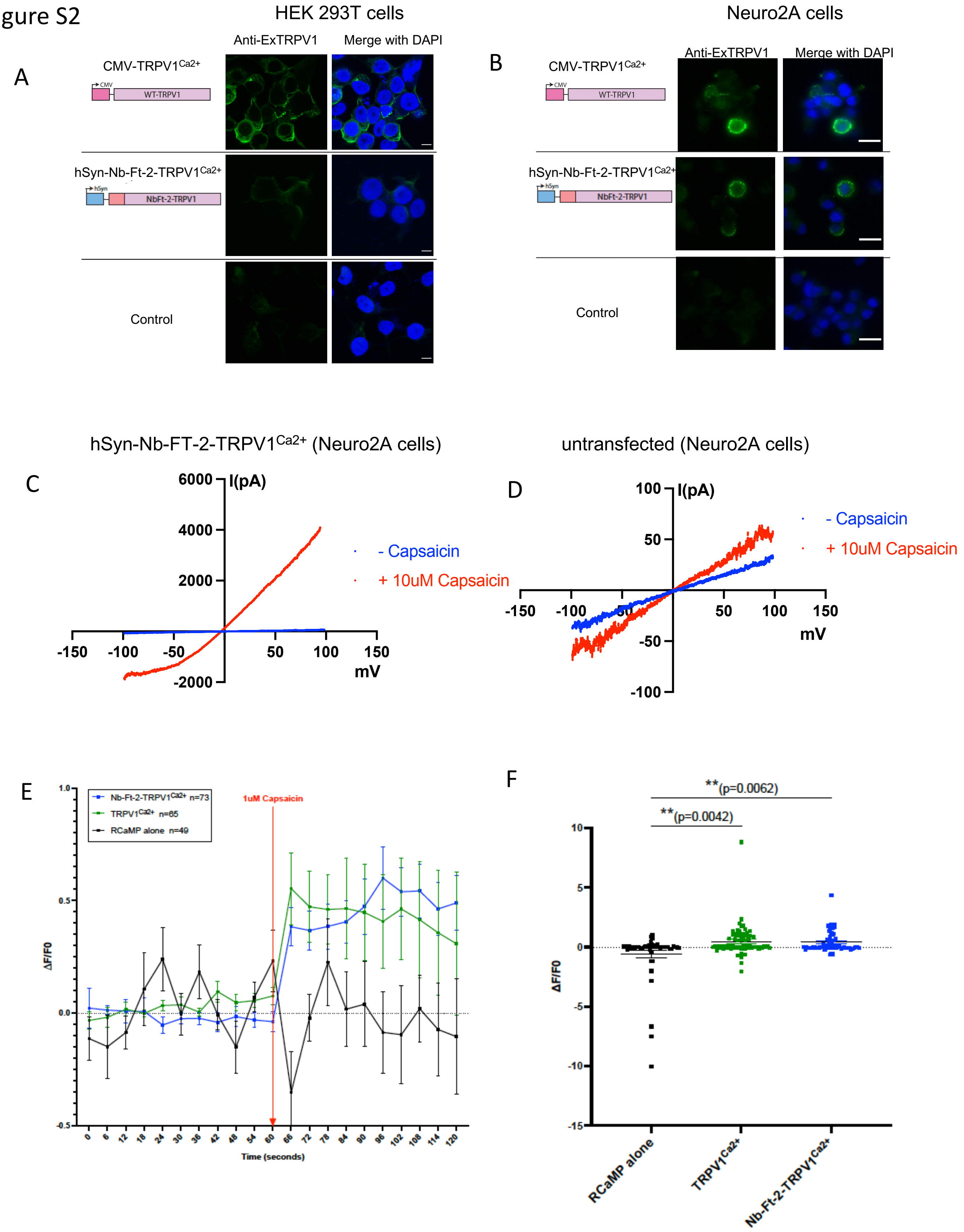

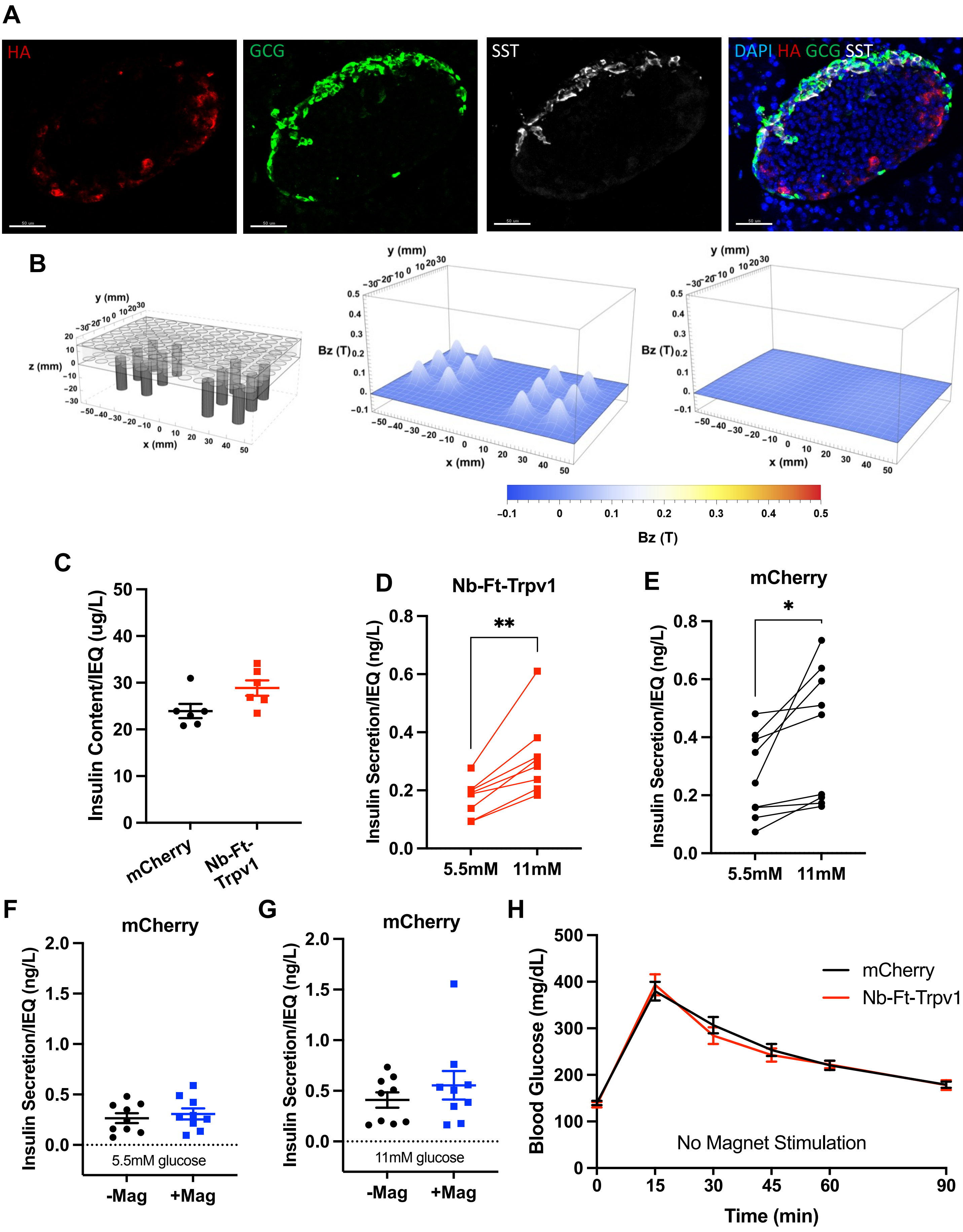
